# Multiframe Evolving Dynamic Functional Connectivity (EVOdFNC): A Method for Constructing and Investigating Functional Brain Motifs

**DOI:** 10.1101/2021.05.18.444678

**Authors:** Robyn L. Miller, Victor M. Vergara, Godfrey Pearlson, Vince D. Calhoun

## Abstract

The study of brain network connectivity as a time-varying property began relatively recently and to date has remained primarily concerned with capturing a handful of discrete static states that characterize connectivity as measured on a timescale shorter than that of the full scan. Capturing group-level representations of temporally evolving patterns of connectivity is a challenging and important next step in fully leveraging the information available in large resting state functional magnetic resonance imaging (rs-fMRI) studies. We introduce a flexible, extensible data-driven framework for the stable identification of group-level multiframe (movie-style) dynamic functional network connectivity (dFNC) states. Our approach employs uniform manifold approximation and embedding (UMAP) to produce a continuity-preserving planar embedding of high-dimensional time-varying measurements of whole-brain functional network connectivity. Planar linear exemplars summarizing dominant dynamic trends across the population are computed from local linear approximations to the 2D embedded trajectories. A high-dimensional representation of each 2D exemplar segment is obtained by averaging the dFNC observations corresponding to the n planar nearest neighbors of τ evenly spaced points along the 2D line segment representation (where n is the UMAP number-of-neighbors parameter and τ is the temporal duration of trajectory segments being approximated). Each of the 2D exemplars thus “lifts” to a multiframe high-dimensional dFNC trajectory of length τ. The collection of high-dimensional temporally evolving dFNC representations (EVOdFNCs) derived in this manner are employed as dynamic basis objects with which to characterize observed high-dimensional dFNC trajectories, which are then expressed as weighted combination of these basis objects. Our approach yields new insights into anomalous patterns of fluidly varying whole brain connectivity that are significantly associated with schizophrenia as a broad diagnosis as well as with certain symptoms of this serious disorder. Importantly, we show that relative to conventional hidden Markov modeling with single-frame unvarying dFNC summary states, EVOdFNCs are more sensitive to positive symptoms of schizophrenia including hallucinations and delusions, suggesting a more dynamic characterization is needed to help illuminate such a complex brain disorder.

## 1. INTRODUCTION

The investigation of functional brain network connectivity (FNC) as a time-varying property in resting state functional magnetic resonance imaging (rs-fMRI) studies began relatively recently, and to date has remained primarily concerned with capturing a handful of discrete static states that characterize connectivity as measured on a timescale shorter than that of the full scan [1–35]. Temporal variation in fMRI has been employed primarily to establish evidence of stable hemodynamic covariation between pairs of functionally or anatomically-defined brain regions or functionally coherent distributed spatial networks. Although initially controversial [36–39], research extending this paradigm from so-called “static” scan-length patterns of functional integration into the analysis of transient but replicable patterns of covariation between functional networks has gained a strong foothold in recent years [40–46]. A substantial amount of this work however focuses on the separation of windowed, time-resolved, connectivity measures into patterns that are consistently transiently evident across subjects. Typically, the dynamics are then treated as a discrete (memoryless) Markov process, characterized by the probability of transitioning from any one of the summary states at time *t* to the same or another at time *t* + 1. The simplifying assumptions that (1) a small number of snapshot summary connectivity patterns capture the functionally important variations large scale brain connectivity on shorter timescales and that furthermore (2) brain dynamics are Markovian, are useful starting points, but stop short of revealing how complex, fluidly varying reconfigurations of whole brain connectivity reflect the myriad dimensions of brain health and dysfunction researchers seek to understand. Capturing group-level representations of temporally evolving patterns of connectivity is a challenging and important next step in fully leveraging the information available in large resting state functional magnetic resonance imaging (rs-fMRI) studies. We introduce a flexible, extensible data-driven framework for the identification of group-level multiframe (movie-style) dynamic functional network connectivity (dFNC) states. Our approach employs uniform manifold approximation and embedding (UMAP) to produce a planar embedding of the high-dimensional whole-brain connectivity dynamics that preserves important characteristics, such as trajectory continuity, of the dynamics in the native high dimensional state space. The method is shown to produce naturalistic, interpretable, evolving dynamic functional network connectivity trajectory motifs (EVOdFNCs) whose role in the dynamic connectomes of schizophrenia patients and healthy controls differ significantly and interpretably, suggesting that methods such as ours that holds promise for identifying more sophisticated dynamical biomarkers from resting-state fMRI than has thus far been possible.

## 2. METHODS

### 2.1 Data

We use data from a large, multi-site eyes-open resting-state fMRI study with approximately equal numbers of schizophrenia patients (SZs) and healthy controls (HCs) (*n*=311, nSZ=150). Imaging data for six of the seven sites was collected on a 3T Siemens Tim Trio System and on a 3T General Electric Discovery MR750 scanner at one site. Resting state fMRI scans were acquired using a standard gradient-echo echo planar imaging paradigm: FOV of 220 × 220 mm (64 × 64 matrix), TR = 2s, TE = 30ms, FA = 770, 162 volumes, 32 sequential ascending axial slices of 4 mm thickness and 1 mm skip. Subjects had their eyes closed during the resting state scan. The data was preprocessed with a standard pipeline and decomposed with group independent component analysis (GICA) into 100 group-level functional network spatial maps with corresponding subject-specific timecourses. Through a combination of automated and manual pruning, *N*=47 functionally identifiable resting state networks (RSNs) were retained (**Figure 1**). The remaining network timecourses were detrended, despiked and orthogonalized with respect to estimated subject motion parameters. Subject specific spatial maps (SMs) and timecourses (TCs) were obtained from the group level spatial maps via spatio-temporal regression. The timecourses were detrended, despiked and subjected to some additional postprocessing steps (details in [35]). All subjects in the study signed informed consent forms. Symptom scores for patients were determined using the Positive and Negative Syndrome Scale (PANSS) [47]. The focus in this paper will be on the six symptoms (Delusions, Grandiosity, Hallucinations, Suspiciousness/Persecution, Preoccupation, Unusual Thought) that load on the “Positive Factor” in a heavily used factor analysis study of PANSS symptoms [48].

**Figure 1.**
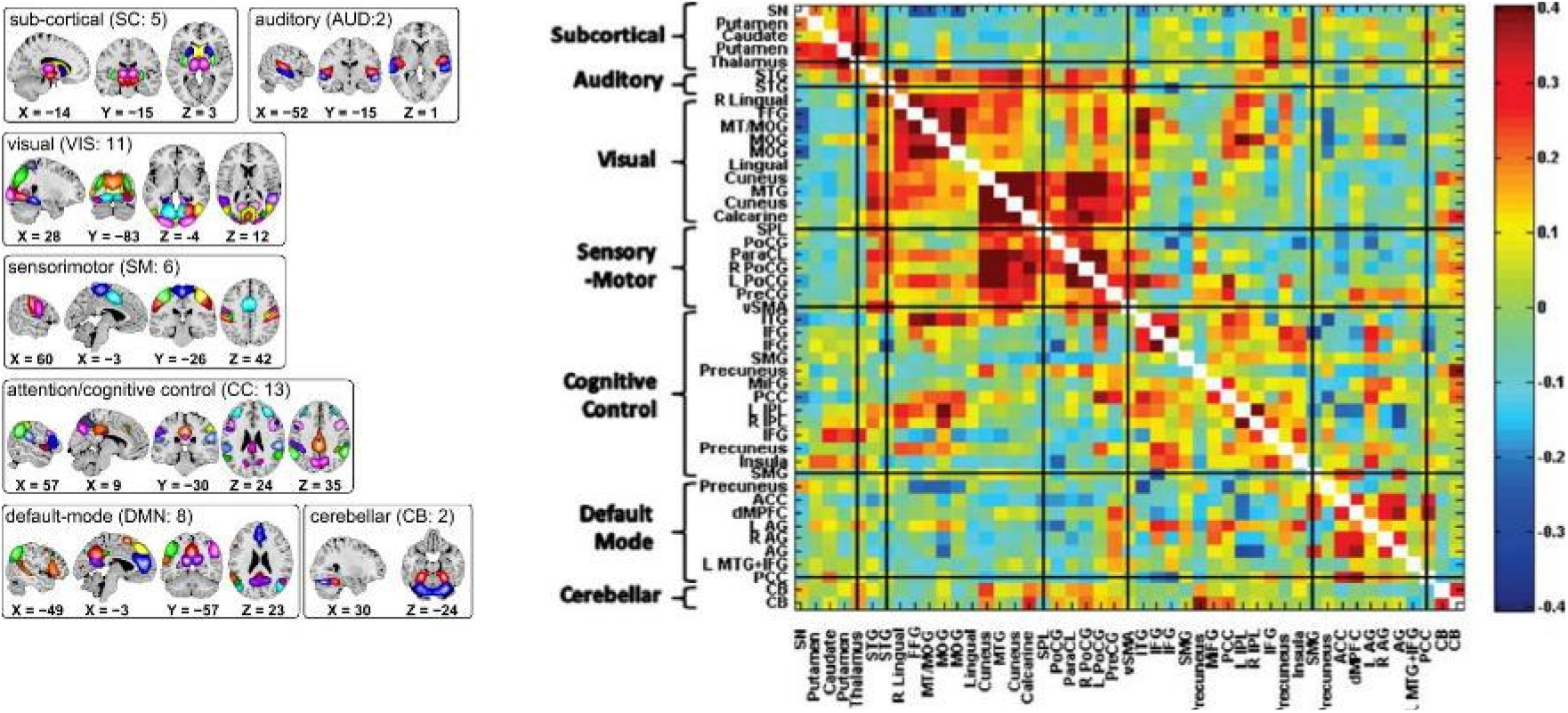
(Left) Composite maps of the 47 resting state networks use in this study, organized according to functional domain with each network in the indicated domain shown in a different color [Damaraju et al, 2014]; (Right) Population means of pairwise correlations between RSN timeseries; The order in which networks and functional domains (shown along the y-axis) are presented here is consistent through all figures in this paper.

### 2.2 Dynamic Functional Network Connectivity

Dynamic functional connectivity (dFNC) between RSN timecourses was estimated using sliding window correlations. Following protocols from published studies on dynamic connectivity [35], we employed a tapered rectangular window length of 22 TRs (44 seconds), advanced 1 TR at each step, and computed pairwise correlations between RSN timecourses within these windows. After dropping the first 3 and final TRs, this procedure yields a 47(47 – 1)/2 = 1081-dimensional dFNC measure on each of 136 windows of length 22TRs for each subject. Clustering this collection of time-resolved connectivity observations using Matlab’s implementation of k-means clustering (Euclidean distance, 2000 iterations, 250 repetitions, 5 clusters chosen according the elbow criterion) produces five non-varying cluster centroids, often referred to as “dynamic states” or dFNC states (**Figure 2**) reported in previous studies [35, 49–58]. We mention them because they are referenced later in the results section of this paper. To distinguish time-blind non-varying states from EVOdFNCs, we will refer to them as nonvarying “snapshot” dFNC states (SNAPdFNC). There are two modularized SNAPdFNC states dominated by strong positive auditory-visual-sensorimotor (AVSM) connectivity: one (State 1) has strong negative connectivity between the default mode network (DMN) and the AVSM networks; the other (State 2) presents weak DMN-to-AVSM connectivity. Healthy controls spend more time in both of these AVSM-dominant modularized states than do patients. A third modularized state (State 3) is defined by a contrast between strong negative connectivity between the DMN and the rest of the brain, with diffuse positive connectivity in all other blocks of the connectome. We will use the shorthand, *DMNneg*, in future references to this particular modularized SNAPdFNC state which is more occupied by patients than controls. The other two SNAPdFNC states (states 4 and 5) show much less modularity: one (state 4) is diffusely hyperconnected and more occupied by controls; the other (state 5) is diffusely disconnected and more occupied by patients [35].

**Figure 2.**
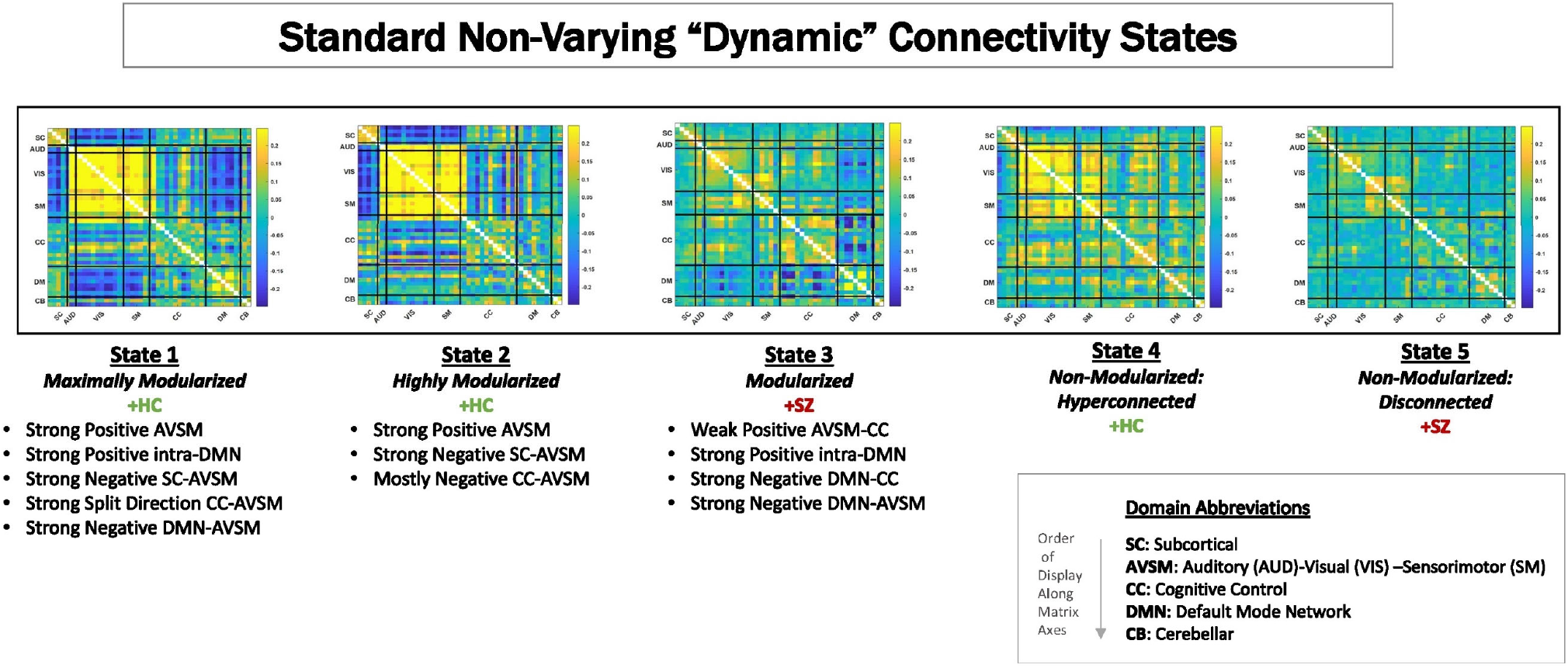
Centroids from clustering time-resolved connectivity measures without in a time-blind manner. These are called “dFNC states”. For this data, as has been shown in previous publications (Damaraju et al, 2014; Miller et al., 2016; Yaesoubi et al., 2017; Espinoza et al., 2019; Rashid et al., 2019), there are five characteristic states that go from maximally modularized to weakly connected. Studies consistently show that schizophrenia patients spend more time in States 3 and 5 on average than do controls, while controls spend more time on average than patients in States 1, 2 and 4.

### 2.3 Planar Embedding

We apply a Matlab implementation [59, 60] of uniform manifold approximation and embedding (*UMAP*) to embed all 1081-dimensional dFNCs into the plane. While both UMAP and t-distributed stochastic neighbor embedding (TSNE) preserve high-dimensional local structure in the lower dimensional embedding, UMAP holds onto more global structure than TSNE and is also significantly more efficient.

Key user-chosen parameters in UMAP are the *number of nearest-neighbors* (n_neighbors) around which the high-dimensional local metrics are defined, and the *minimum distance* (min_dist) which parameterizes proximity in the low dimensional embedding. Since dFNCs are computed on sliding windows that advance one TR at a time, they exhibit considerable temporal smoothness in their native high-dimensional space (mean elementwise squared distance between successive dFNC measures is less than 0.0006). The high-dimensional trajectories are also somewhat “densely packed”, i.e. for any fixed dFNC observation, the average mean squared elementwise distance between that observation and its 10,000 nearest neighbors *from different subjects* is less than 0.01. The ideal embedding for our purposes would preserve intrasubject trajectory continuity and inter-subject trajectory proximity. This within-subject smoothness is an intrinsic feature of the actual dynamics, which we experimentally optimized UMAP’s two main parameters (n_neighbors=25; mindist=0.75) toward conserving in the planar embedding (**Figure 22** (Supplementary)). It is worth quickly noting that linear dimension-reduction methods such as PCA or ICA produced diffuse 2D clouds lacking intra-subject temporal continuity (**Figure 3**), while t-SNE performed more than 200-fold more slowly than UMAP on this data, and thus was not practical for achieving a planar embedding of the entire dataset.

**Figure 3.**
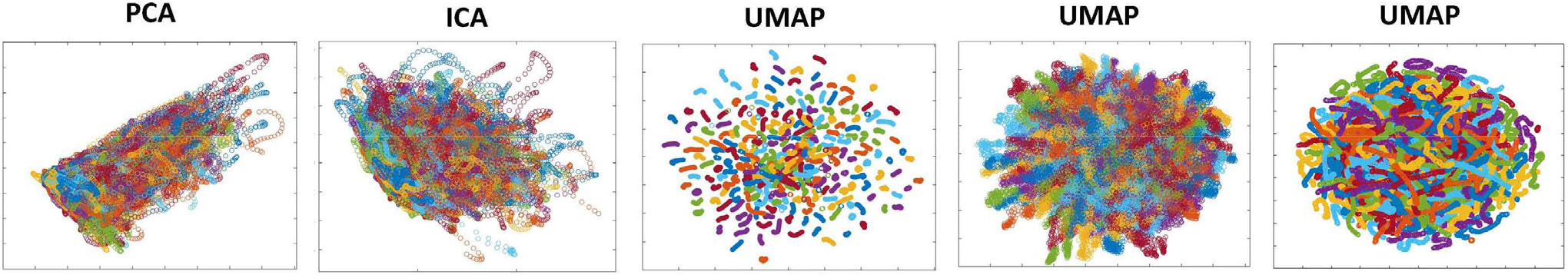
Applying different data reduction methods to the 1081-dimensional dFNC observations yields various planar presentations. PCA (column 1) and spatial Independent Component Analysis (ICA) (column 2) are “cloudy” and do not produce continuous, well defined, subject-level trajectories; UMAP with n_neighbors=200, min_dist=0.1 (column 3) has a large periphery of isolated geometrically compressed trajectories; UMAP with n_neighbors=200, min_dist=0.8 (column 4) results in a crowded but diffuse embedding that blurs within-subject temporal trajectory continuity. UMAP with the parameters used in this study, n_neighbors=25 and min_dist=0.75, embeds individual subject trajectories as continuous curves which are densely intermingled at a group level, qualitatively reflecting the high-dimensional reality. In all panels, points corresponding to each subject are plotted in the same color.

Since the UMAP embeddings are not fully deterministic, we run the algorithm *R* = 25 times on the input dFNC data 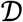 and use the average 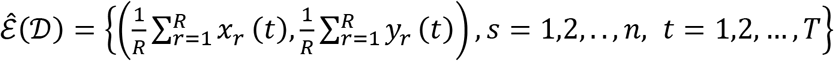, of the *R* individual 2D embeddings 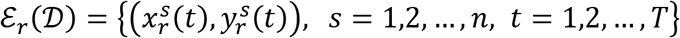. This averaging procedure stabilizes the embedding but also de-densifies the group level geometry (**Figure 23** (Supplementary)), reducing its qualitative fidelity to the native high-dimensional setting.

### 2.4 Linearized 2D dFNC Trajectory Segments and Gradient Exemplars

Due to our choice of UMAP parameters within intervals that preserve intra-subject trajectory continuity, the majority of each subject’s high-dimensional dFNC trajectory Γ(*t*) = (*υ*_1_(*t*), *υ*_2_(*t*),…,*υ*_1081_(*t*)) embeds, under the *r^th^* run of UMAP, into an approximately continuous segment *γ*_r_(*t*) = (*x_r_*(*t*), *y_r_*(*t*)) in 2D (**Figure 4**). Continuity is preserved under summation, so the average 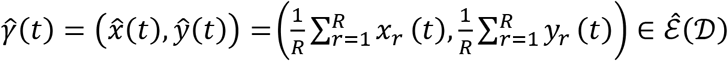 will also be continuous. We collect all continuous trajectory sub-segments (CTSs) 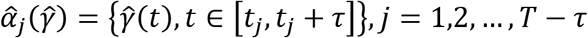 along 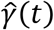 of temporal duration τ = 44 (twice the window length used to estimate the dFNC observations in 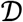). Each 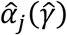 can be approximated by a line *L_j_*, yielding a reduced characterization 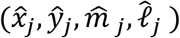 of 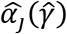 in terms of its length 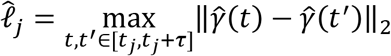, its geometric midpoint 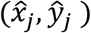 and the slope 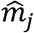 of its linearization *L_j_* (**Figure 5**). The linearized trajectory segment (LTS) triples: 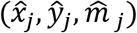 are then clustered (**Figure 15**) with k-means (Matlab implementation, squared Euclidean distance, 2000 iterations, 250 repetitions) where the number of clusters, *K =* 10, was chosen according to the elbow-criterion. From each of the *K* = 10 LTS cluster centroids: 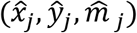 we induce a 2D line segment of length *d_i_* (equal to the mean length of all LTSs in that cluster) and slope 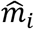 centered at 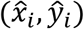 (**Figure 4**, **Figure 5**). These segments are, roughly speaking, *gradients* of the CTSs, which are in turn averages of continuous embedded segments of the high dimensional dFNC dynamics. The collection of segments induced by LTS cluster centroids will be called *linear trajectory exemplars* or just *exemplars*.

**Figure 4.**
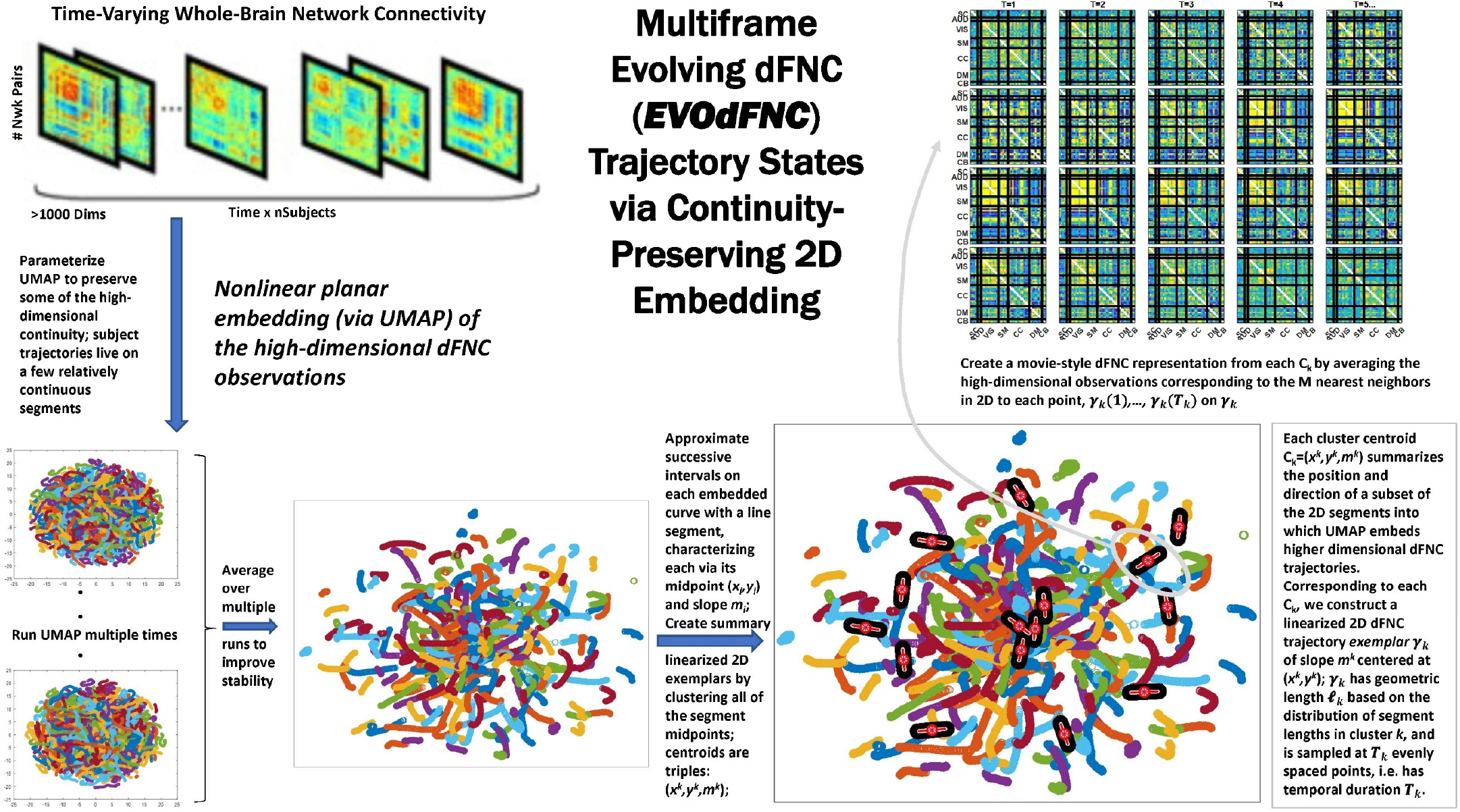
Overview schematic exhibiting the pipeline for producing representative evolving multiframe high-dimensional states of dynamic functional network connectivity (EVOdFNCs). High-dimensional dynamic functional connectivity assessed as pairwise correlations on sliding windows through each subject’s GICA network timecourses (top left) are embedded into 2D using uniform manifold approximation and projection (UMAP). The embedding is done *R* times (bottom leftmost), then averaged over the *R* runs to improve stability (bottom second from the left). Continuous trajectories in the averaged embedding are treated as locally (i.e., sub-segments of temporal duration τ = 44) linear. Directional and positional trends in the 2D dynamics are then captured by clustering the midpoints and slopes of these local linearizations, resulting in a set of summary tangents to the embedded trajectories, *linearized trajectory exemplars* (bottom right). The exemplars are then “inverted” back to the native dFNC dimensionality, as EVOdFNCs, by associating to each of τ evenly points along an exemplar, the average of the high-dimensional dFNCs that map to that point’s *n* nearest neighbors in 2D (top right).

**Figure 5.**
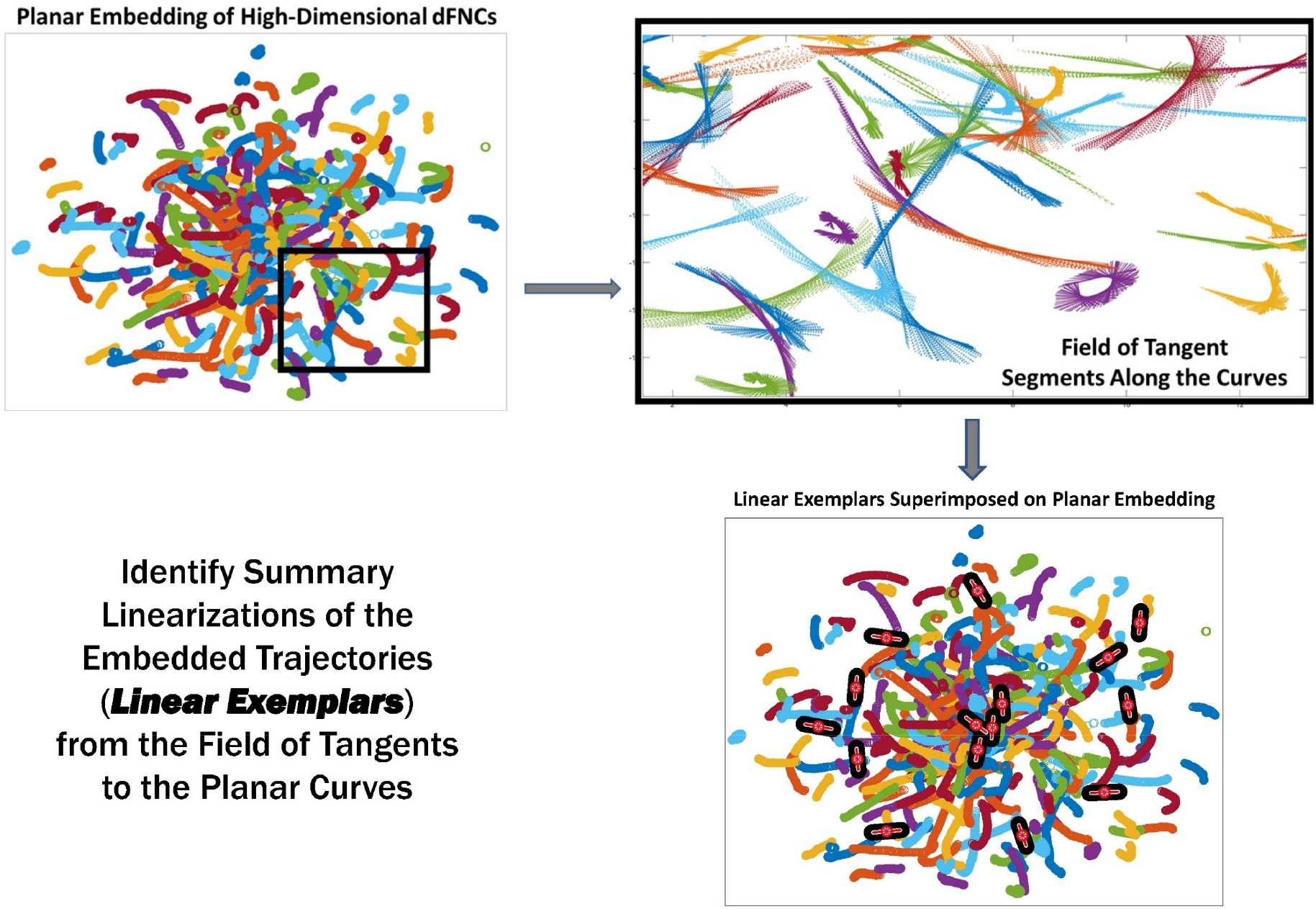
The linear trajectory segments (LTSs) (top right) along the boxed embedded trajectories (top left). These local linearizations are clustered to identify a set of linear trajectory exemplars (bottom right, superimposed black and red segments) capturing localized directional trends in the dynamics

**Figure 6.**
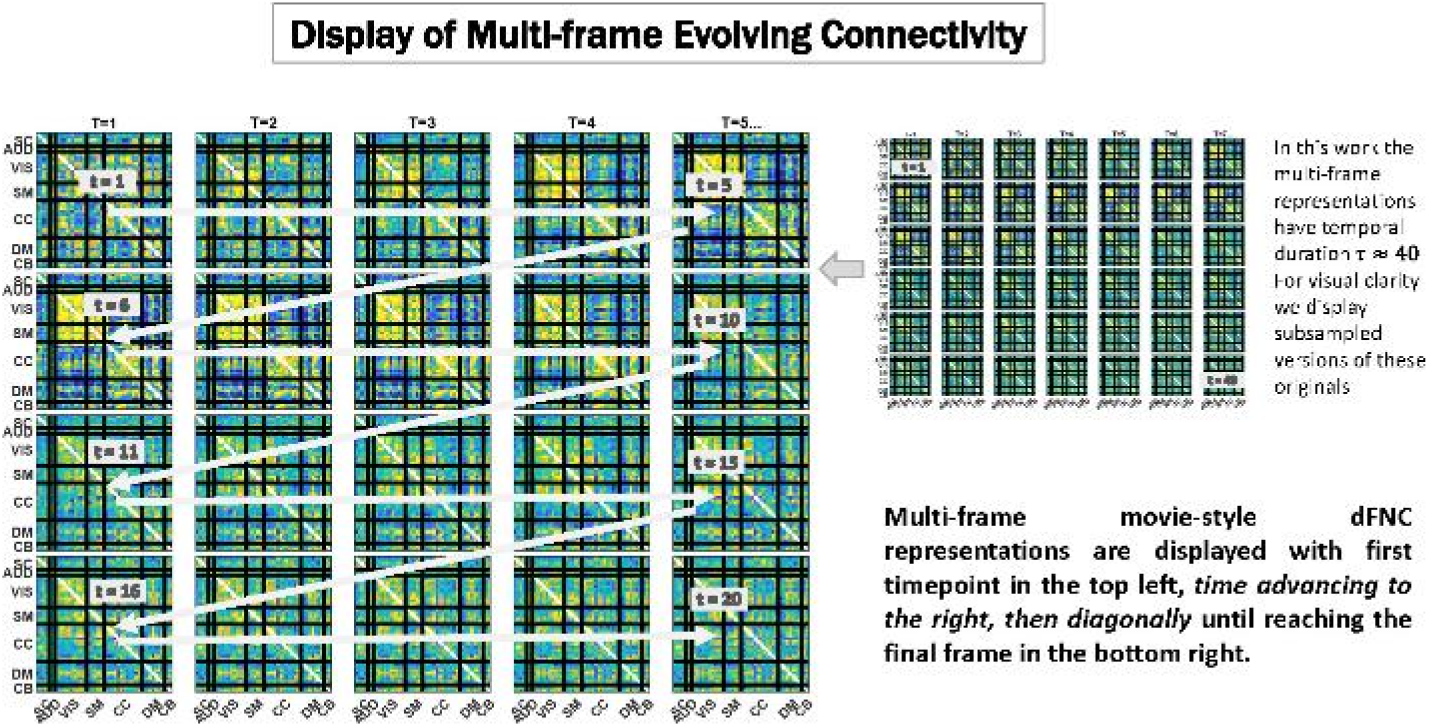
We display temporally evolving dFNCs as a sequence of frames in which the first timepoint is always the top leftmost subplot, and with time advancing from left to right, then clockwise diagonally back to the first subplot in the subsequent row, until reaching the final frame of the sequence shown in the bottom rightmost subplot. Since these are very dense displays, the EVOdFNCs in this work, which have duration 44 timepoints are subsampled to show only every other frame in figures below. To avoid more crowding, colorbars are not displayed, but the range is fixed across EVOdFNCs, centered at 0 and bounded in [–*q, q*] where *q* denotes the 95*^th^* percentile of all magnitudes in the collection of EVOdFNCs.

### 2.5 Prototype High-Dimensional Evolving dFNC Basis States (EVOdFNCs)

Because UMAP is not straightforwardly invertible, the 2D linear trajectory exemplars above cannot easily be mapped back directly into the native dimension of the dFNC space. To obtain the high dimensional dFNC trajectory segment *γ* of integer temporal duration τ corresponding to a given 2D linear trajectory exemplar *υ*, (i.e., the “data driven inverse” of *υ*), we average the high-dimensional observations corresponding to the *n* = 25 nearest 2D neighbors of each of τ evenly spaced points along *υ* (**Figure 7**). The number of 2D neighbors used to invert UMAP, *n* = 25, was chosen to match the number of nearest neighbor parameters employed in the UMAP embedding. This operation effectively “lifts” a localized 2D linear trajectory exemplar back into high-dimensional dFNC space. The 2D linear trajectory exemplars are, by construction, concentrated in more densely occupied parts of the plane, and the continuity preserving parameterization of UMAP encourages the high-dimensional data-driven inverse of each 2D linear trajectory exemplar to exhibit naturalistic smoothness (**Figure 10**). These high dimensional inverse images of the 2D linear trajectory exemplars are each multiframe representations of evolving functional network connectivity (EVOdFNCs).

**Figure 7.**
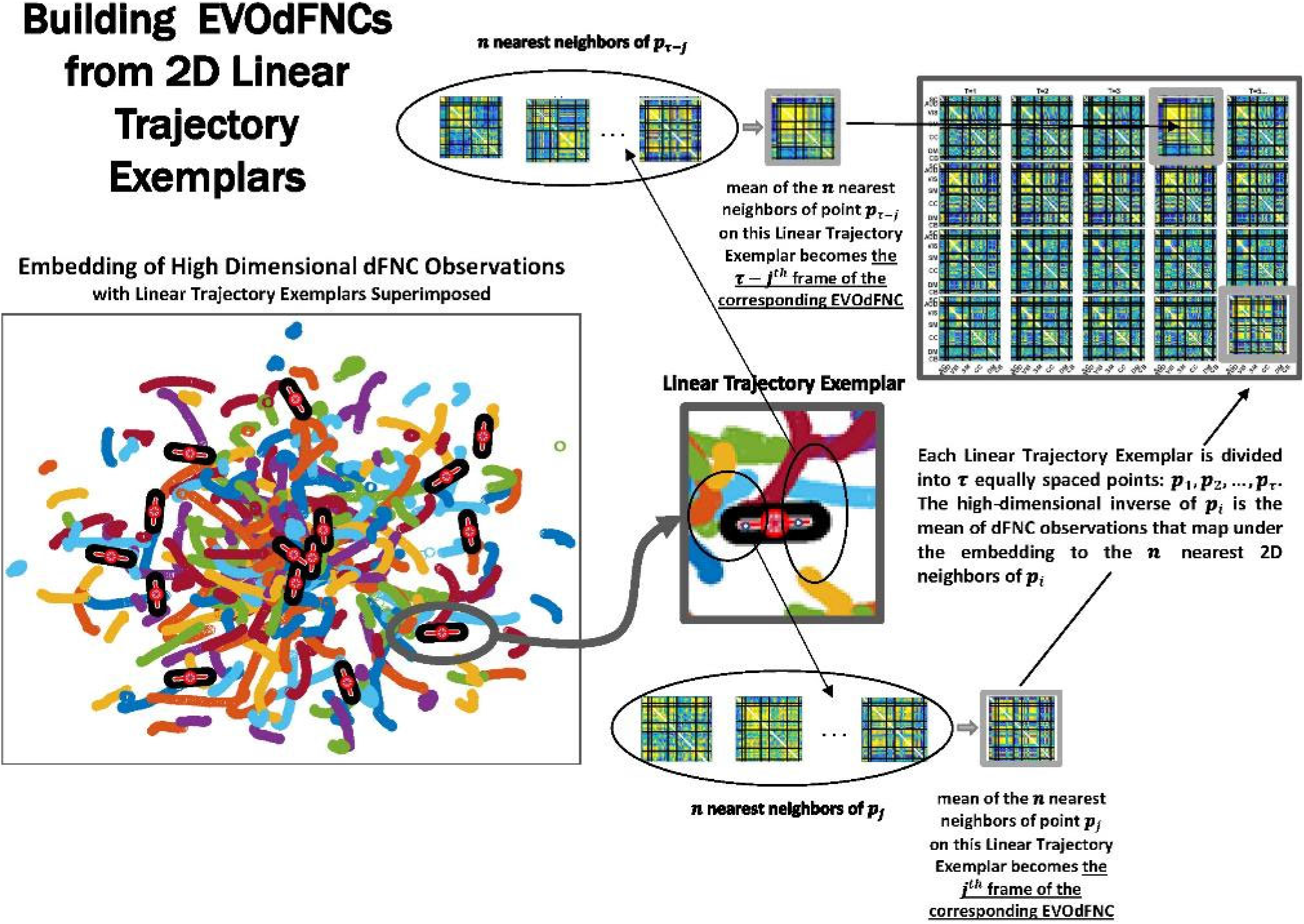
Exemplars are divided into τ evenly-spaced points: *p*_1_, *p*_2_,…,*p*_τ_ (where τ is the mean temporal duration of the piecewise linear approximations of continuous trajectories that are inputs to the clustering). The geometric length of each exemplar is the based on the lengths of the continuous trajectory segments belonging to the cluster they represent. Timepoint *t* = *i*, *i* = 1,2,…, τ of the multifrane EVOdFNC corresponding to linear exemplar *k* is the average of the observed high-dimensional dFNCs corresponding to the *n* = 25 nearest 2D neighbors of *p_i_*.

### 2.6 Expressing Observed High-Dimensional dFNC Trajectories in Terms of EVOdFNC Basis States

The high-dimensional EVOdFNCs obtained from 2D exemplar linear trajectories are more properly viewed as representations of dominant directional trends than as a hard segmentation of the observed dynamics. We therefore employ them as basis objects through which to parameterize observed high-dimensional evolving connectivity dynamics. This is done by correlating successive timepoints (i.e., *t* = 1,2,…, τ) of each length-τ window *w_i_* in a subject’s observed dFNC sequence with the corresponding set of *t^th^* timepoints from the *K* length-τ EVOdFNCs (**Figure 8**). For each *w_i_* and each *t* ∈ {1,2, …, τ}, we identify the EVOdFNC *k* ∈ {1,2, …, *K*} whose *t^th^* timepoint has the highest correlation with the *t^th^* timepoint of observed dFNC window *w_i_*, yielding a length-τ vector 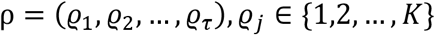.

**Figure 8.**
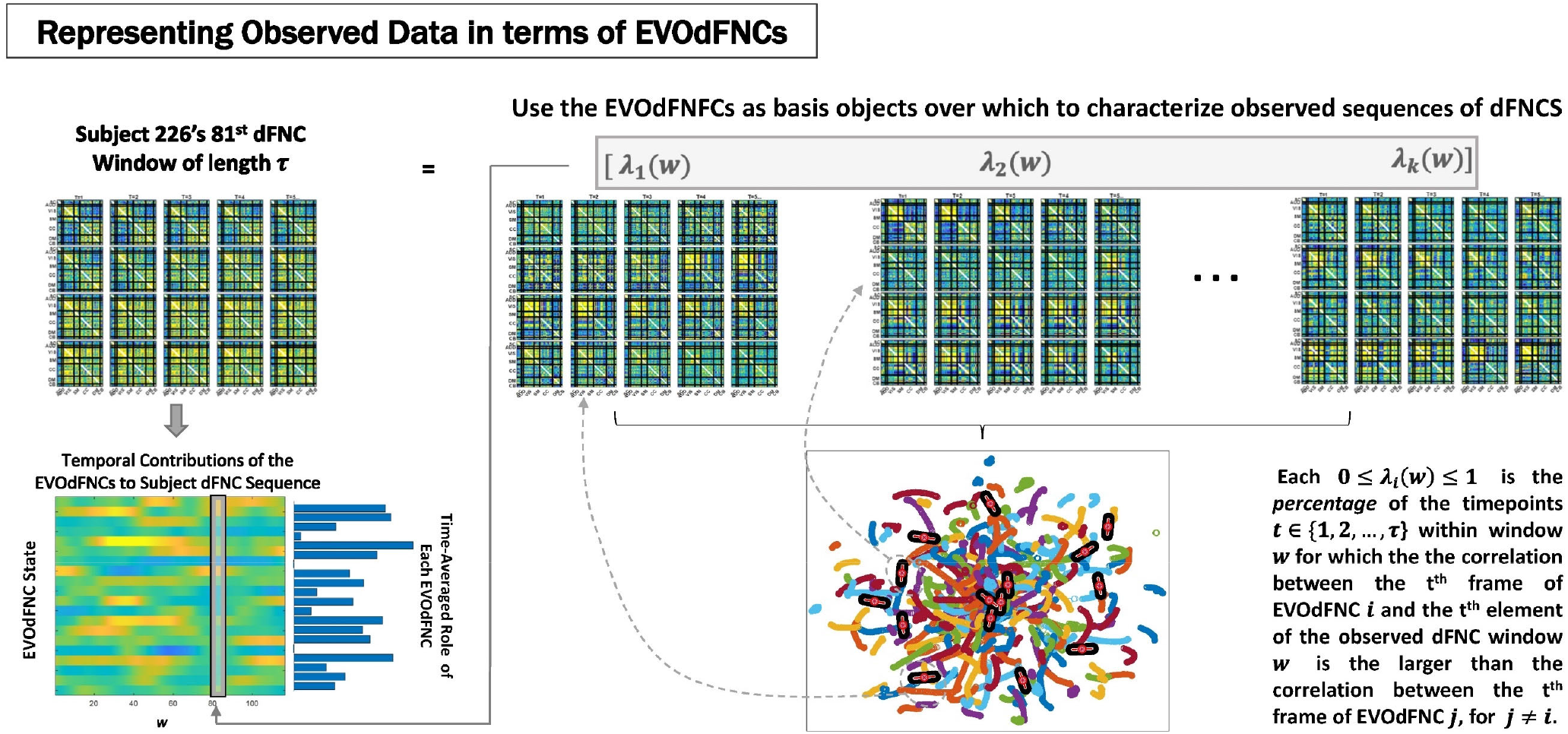
Represent each length-τ window *w* from the observed subject dFNCs as a weighted combinations of the *K* EVOdFNC states (top row); Rather than simply regressing full 1081τ = 47,564-dimensional dFNC windows on the *K* 1081τ-dimensional EVOdFNCs, which ends up highlighting the relationship, we go timepointwise through the data window and EVOdFNCs, expressing each timepoint *t* of the data in terms of the corresponding timepoint in each EVOdFNC, then averaging this evolving correspondence over the window *w* between the observation and each EVOdFNC *k* to obtain the weight *β_k_*(*w*) for the *k^th^* EVOdFNC on the *r^th^* length-τ window of the subject’s dFNC sequence. This yields a K-variate timeseries expressing the subject’s observed sequence of dFNCs in terms of time-indexed weightings on the EVOdFNCs (bottom left).

The weight *λ_k_*(*w_i_*) ∈[0,1] on each *k* ∈ {1,2, …, *K*} is the proportion of timepoints 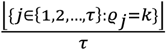 in *w_i_* that most resemble (are most correlated with) the same timepoint in EVOdFNC *k*. This approach both normalizes the weights in [0,1] and accommodates the possibility that within any given time window of length τ, a subject’s dFNC sequence might contain subintervals that are best represented by the corresponding subinterval of different EVOdFNCs. From this, we obtain a *K*-variate timeseries of EVOdFNC weights 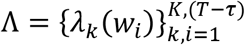 for each subject (**Figure 8**, bottom left), capturing the relative *representational importance* (RI) of each EVOdFNCs to successive length-τ intervals of the subject’s observed high-dimensional dFNC trajectory.

### 2.7 Obtaining *meta*-EVOdFNCs from Multivariate EVOdFNC Timeseries

Since EVOdFNCs are inverse images of 2D segments in a planar embedding of very high-dimensional data, using weighted combinations of the EVOdFNCs (as opposed, e.g., to a binary “occupancy” approach) is arguably a better strategy for capturing the evolving variability of high-dimensional dFNC observations. Toward this end, we cluster the time-indexed weight vectors from the multivariate EVOdFNC timeseries into *M* = 10 clusters (elbow criterion, Matlab’s k-means implementation, squared Euclidean metric, 2000 iterations, 500 repetitions). We then induce *M meta*-EVOdFNCs as centroid-weighted sums of the basis EVOdFNCs (**Figure 9**).

**Figure 9.**
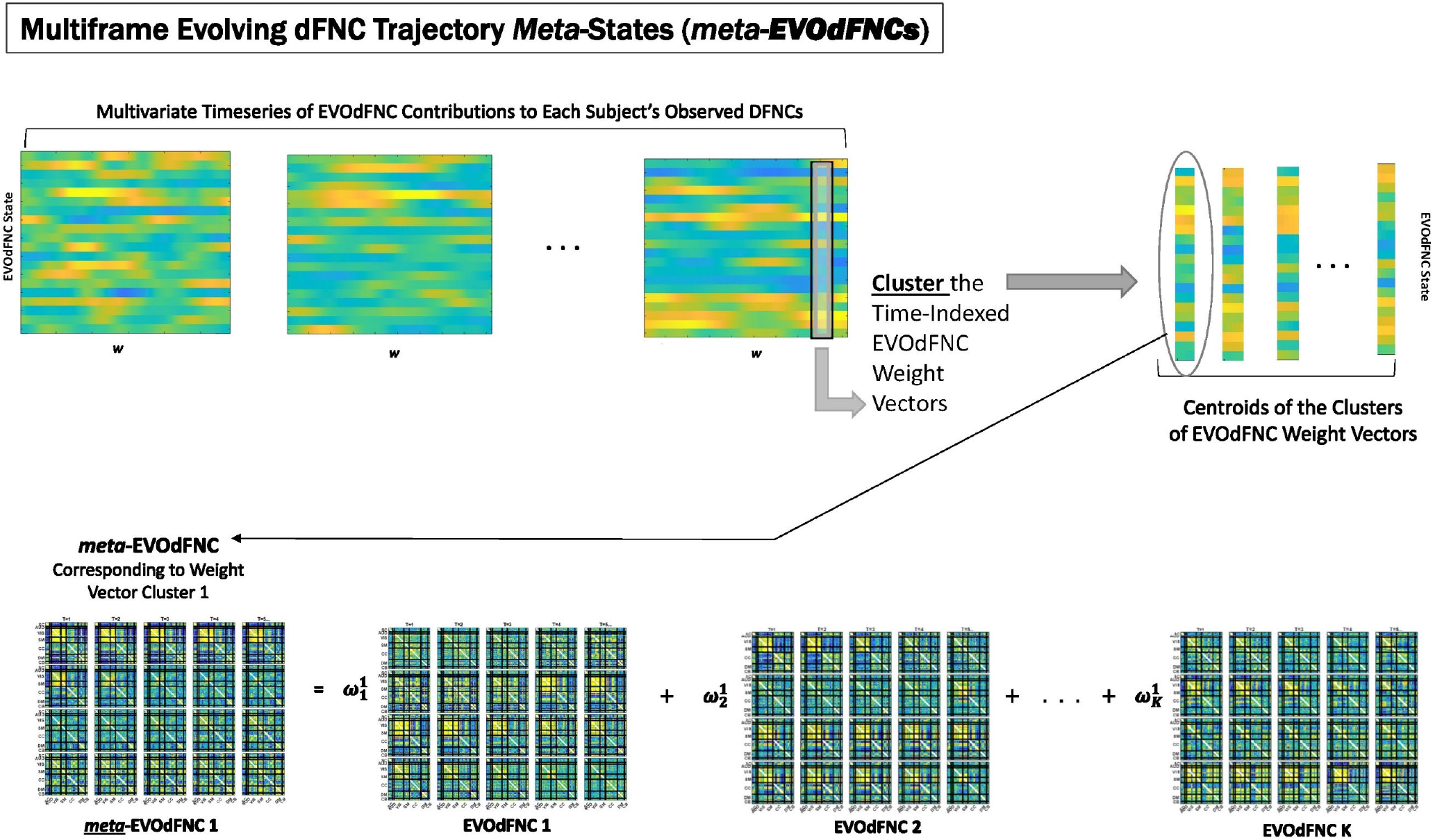
EVOdFNCs are high-dimensional data-driven inverses of exemplar 2D linear trajectories that are relatively sparse in the space of linearized 2D trajectory segments. A fuller range of the structure and dynamics exhibited by actual high-dimensional dFNC observations can be captured by expressing each length-τ observed dFNC sequence as a weighted combination of the EVOdFNCs. From the multivariate EVOdFNC timeseries we induce a collection of *meta*-EVOdFNCs by clustering the time-indexed weight vectors (top row) and applying the weight-vector cluster centroids to the EVOdFNC basis states (bottom row).

### 2.8 Statistical Modeling

All reported SZ effects are from a multiple regression on gender, age, head motion (mean frame displacement) and SZ. Reported positive PANSS symptom score effects are from a multiple regression on the six positive PANSS symptoms (Delusions, Grandiosity, Hallucinations, Suspiciousness/Persecution, Preoccupation, Unusual Thought) from the Lindenmeyer five-factor PANSS model [48], in addition to age, gender and head motion. The purpose of this model is to identify the effect of each positive symptom corrected for the contributions of the others, in a manner indifferent to, i.e., summing over, the various profiles of negative and general symptoms that subjects might manifest. While negative symptoms in the SZ subjects analyzed here are highly intercorrelated (mean correlation between negative symptom pairs = 0.31 (std. dev = 0.12)), positive symptoms are not (mean correlation between positive symptom pairs = 0.12 (std. dev = 0.13)). Schizophrenia effects are only reported when significant at the *p* < 0.05 level *after* correction for multiple comparisons. All displayed PANSS symptom effects are significant at the *p* < 0.05 level, and remain significant at this level after correction for multiple comparisons where specifically indicted.

## 3. RESULTS

### 1. Representational Importance of EVOdFNCs: Effects of Schizophrenia and Positive Symptoms

We find widespread schizophrenia effects on the representational importance (RI) of each of the 10 EVOdFNCs in subject data (**Figure 10**). The evolving motifs fluidly move through transient states of connectivity that resemble familiar formations obtained from the basic time-blind clustering into single transiently realized connectivity patterns (**Figure 2**). Consistent with published results [35, 49, 53, 54, 57] on occupancy rates of time-blind SNAPdFNC states, we find here that: strongly modularized and hyperconnected patterns feature more prominently in EVOdFNCs with greater representational importance in controls (1, 5, 7 and 8) and in EVOdFNCs whose representational importance in controls is not statistically distinguishable from that in patients (4, 6, 9); weak connectivity and modularized negative DMN-to-other (DMNneg) patterns (2, 3) feature more prominently in EVOdFNCs with significantly higher representational importance in patients. A novel modularized pattern of functional organization, not seen in time-blind SNAPdFNC states, appears in EVOdFNC 10, which features a persistent stretch of strong modularized negative SM-VIS/CC/DMN connectivity. EVOdFNC 10 has significantly greater RI in patients, and its distinctive modularity eventually dissolves into unstructured weak connectivity. EVOdFNC 7, with greater RI in controls, transitions from from a weakly connected negative DMN-to-other structure (more characteristic of patients in time-blind analysis, e.g. SNAPdFNC State 3) into another novel pattern that is strongly modularized with positive DMN-VIS/SM connectivity. EVOdFNC 2 (+SZ, RI) and EVOdFNC 5 (+HC, RI) show roles for disconnected periods leading into and out of modularized structures that are more characteristic of SZ and HC respectively. EVOdFNC 2 (+SZ, RI) also exhibits a transition from something more like SNAPdFNC State 1 (+HC, OCR) to SNAPdFNC State 3 (+SZ, OCR), presenting a path from strong AVSM-focused modularity (+HC, SNAPdFNC OCR) to the DMNneg configuration (+SZ, SNAPdFNC OCR) to weak connectivity (+SZ, SNAPdFNC OCR). EVOdFNCs 1 and 8 (both +HC, RI) show the role of disconnected periods in transitioning between AVSM-centered modularity and hyperconnectivity in healthy people: in EVOdFNC 1 we see modularity dissolving into diffuse dysconnectivity which then uniformly inflates toward diffuse hyperconnectivity, while in EVOdFNC 8, diffuse hyperconnectivity gradually erodes toward greater dysconnectivity away from the AVSM core, then starts manifesting negative DMN/CC-AVSM connectivity from a substrate of dysconnectivity. While EVOdFNCs 2 and 3 both have greater representational importance in patients and both contain patterns similar to SNAPdFNC State 3 (+SZ, OCR), it is the weaker, less dynamically varying EVOdFNC 3 whose RI is significantly further elevated in more delusional patients (**Figure 11**). The RI of EVOdFNCs 4 and 9 are not significantly affected by subject diagnosis, but they reveal patterns of functional organization that do not appear in time-blind dFNC clustering. Specifically, EVOdFNC 4 evolves from strong AVSM-centered modularity (+HC, SNAPdFNC OCR) through a DMNneg configuration (+SZ, SNAPdFNC OCR) connectivity into a modularized pattern in which intra-AVSM and intra-CC connectivity is at approximate parity. EVOdFNC 9 passes through a stage in which intra-CC connectivity is even stronger than intra-AVSM connectivity. Transient periods of whole brain connectivity where connectivity strength within the CC domain equals or exceeds that within AVSM domain block do not appear in time-blind dFNC clustering of this data. EVOdFNCs 4 and 9 both have strong population-wide representational importance (**Figure 19**) indicating that these patterns play a meaningful role in dynamic connectivity despite not appearing in SNAPdFNC clusters centroids.

**Figure 10.**
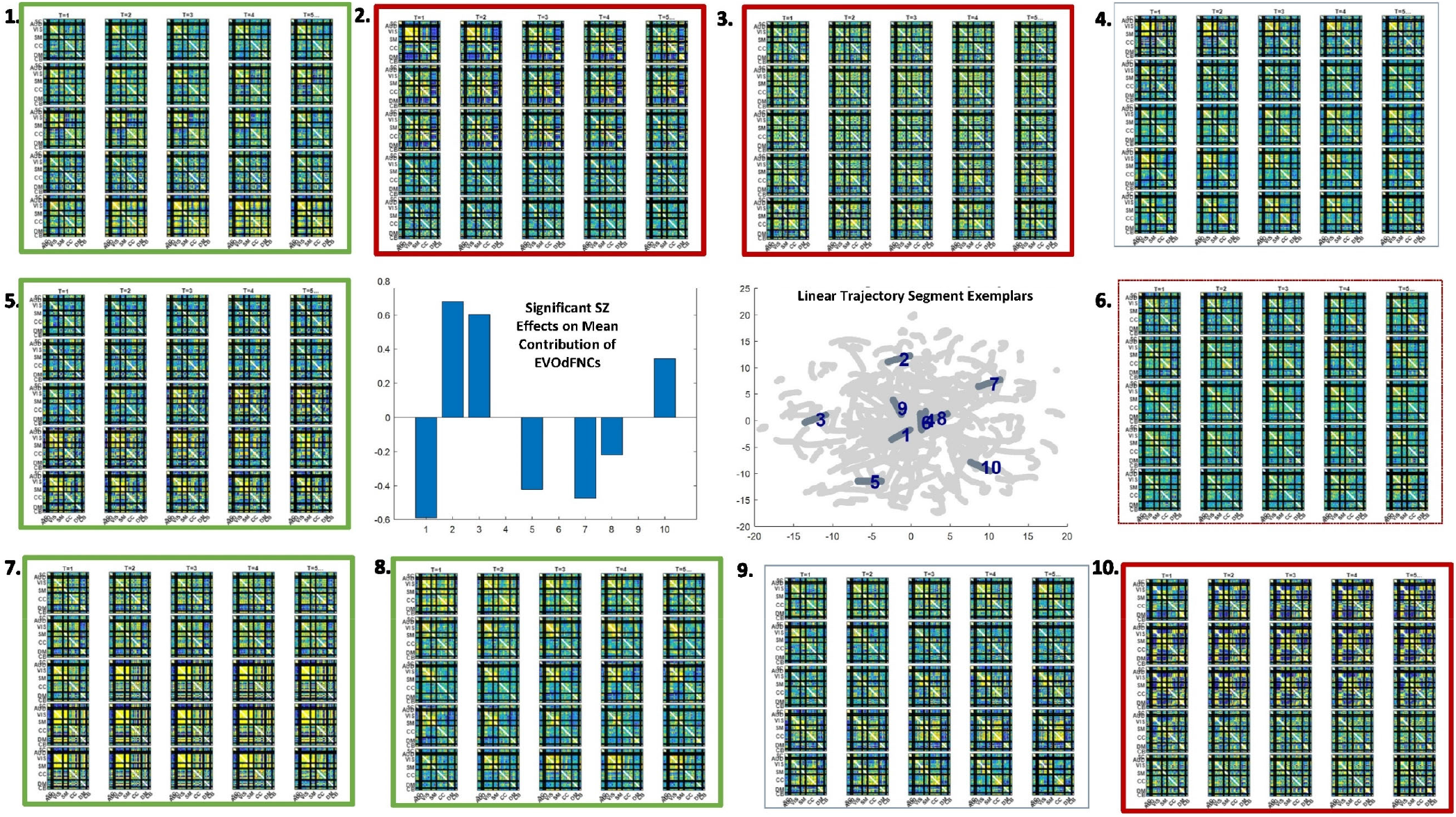
There are pervasive group differences between schizophrenia patients and controls in representational importance of the *K* = 10 EVOdFNCs; thick red (resp. thick green) boxes designate significant positive (resp. negative) association with SZ after correction for multiple comparisons; thin dashed red box designates significant (*p* < 0.025) positive association with SZ that is not significant after correction for multiple comparisons. Omitted colorbar is bounded by [–0.3,0.3].

**Figure 11.**
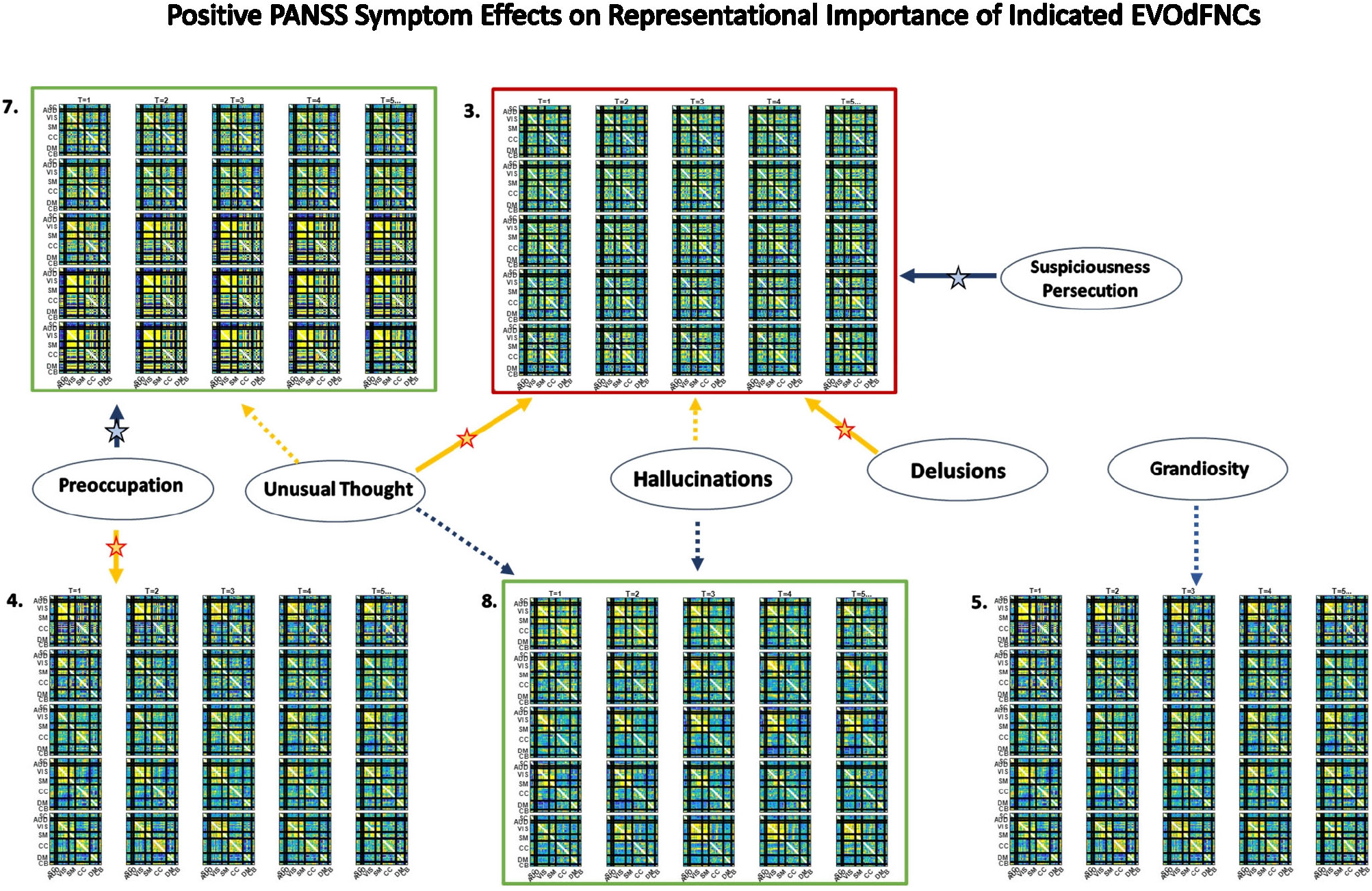
Significant positive symptom effects on the representational importance of EVOdFNCs in high dimensional dFNC dynamics. Thick arrows with stars indicate effects that are significant after correction for multiple comparisons; thin dashed arrows indicate effects that are significant with *p* < 0.05, but not after correction for multiple comparisons. Yellow lines denote positive effects. Blue lines denote negative effects. EVOdFNC 3, the low contrast, low dynamism DMNneg patterned state is a locus of positive symptom effects: RI elevated by Hallucinations, Delusions and Unusual thought, and suppressed by Suspiciousness/Persecution. EVOdFNC 7, dynamically changing from a low contrast DMNneg pattern to an AVSM-dominant hyperconnected pattern is more important in HCs than SZs, but among SZs with high levels of Preoccupation its importance is further significantly suppressed, and elevated in those with high levels of Unusual Thought. Note that positive symptoms exhibit no significant (p<0.05, uncorrected) effects on the occupancy rates of the SNAPdFNCs.

Three of the ten EVOdFNCs start with intervals characterized by modularized DMNneg patterning, i.e., strong negative connectivity between DMN and all other brain areas with positive connectivity everywhere else. The SNAPdFNC state with this pattern is more occupied by SZ patients than HCs. Two of the EVOdFNCs (2 and 3) have higher RI in SZs, one (EVOdFNC 7) in HCs. Two have significant positive PANSS symptom effects (**Figure 11**): one of these (EVOdFNC 3) has higher RI in SZs, the other (EVOdFNC 7) has higher RI in HCs. Although all three EVOdFNCs pass through a DMNneg type of pattern, each embeds this pattern in a different dynamic context (**Figure 12**). EVOdFNC 2 presents a high-contrast DMNneg pattern dynamically dissolving into diffuse dysconnectivity. Its temporal mean (bottom left) resembles that of EVOdFNC 3, which features persistent, low contrast, unvarying DMNneg. Both EVOdFNC 2 and EVOdFNC 3 have higher RI in SZs, but only the lower contrast, less dynamic version, EVOdFNC 3, has significant further relationships to positive symptoms within the patient population (**Figure 11**). The temporal averages of EVOdFNCs 2 and 3 (**Figure 12**, bottom row) are very similar, suggesting that the differential sensitivity of EVOdFNCs 2 and 3 to positive symptoms resides in their temporal patterning. Finally, although in time-blind SNAPdFNC analysis the DMNneg pattern is significantly more occupied by SZs, we find EVOdFNC 7 (**Figure 12**, top right) featuring DMNneg leading into AVSM-dominant, lightly modularized hyperconnectivity to have significantly higher RI in HCs, and also to be sensitive to positive symptoms in SZ patients (**Figure 11**).

**Figure 12.**
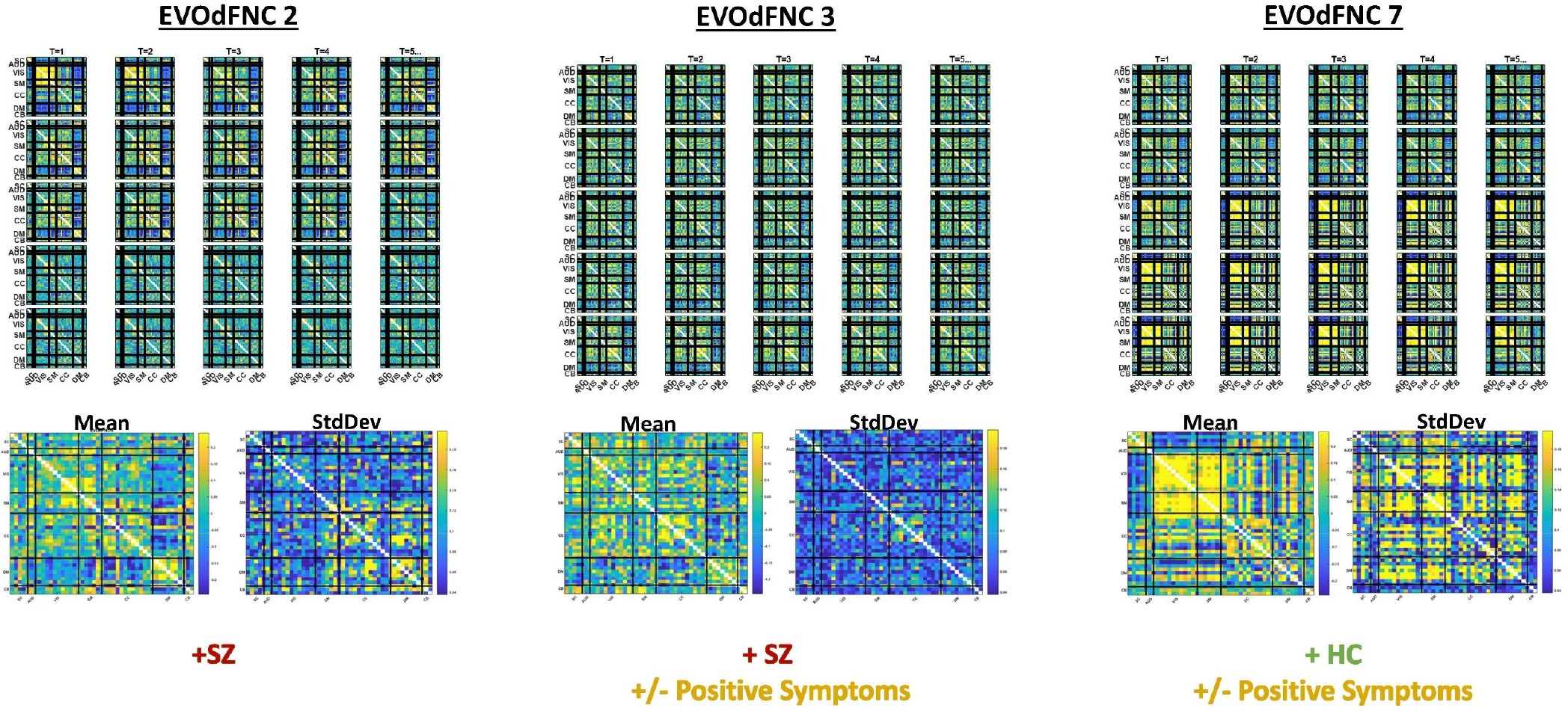
Three EVOdFNCs (2, 3, 7) start with intervals characterized by the modularized DMNneg patterning defined by SNAPdFNC State 3, a state that is more occupied by SZ patients. All three EVOdFNCs start with a DMNneg type of pattern, but each presents a different dynamic context. EVOdFNC 2 (top left) exhibits a high-contrast DMNneg pattern that dissolves into diffuse dysconnectivity. EVOdFNC 3 (top middle) presents a persistent, unvarying low-contrast DMNneg pattern. EVOdFNCs 2 and 3 have very similar temporal means (displayed below each EVOdFNC), with distinguishing characteristics requiring representation of their temporal evolution. Only EVOdFNC 3 has significant positive symptom effects. In EVOdFNC 7, the DMNneg pattern is part of a dynamic progression leading to AVSM-dominant hyperconnectivity. Although DMNneg patterning is significantly associated with SZ in time-blind SNAPdFNC analysis, EVOdFNC 7 has higher RI in HCs, but also is sensitive to positive symptom levels in SZ patients.

### 2. Occupancy Rates of *meta*-EVOdFNCs: Effects of Schizophrenia and Positive Symptoms and Relationship to EVOdFNCs

The 2D linear exemplars that lift to high dimensional EVOdFNCs are relatively sparse in the planar embedding. Many local linearizations of embedded subject dFNC trajectories are not geometrically proximal to an exemplar or are not directionally aligned with the nearest exemplar. Moreover, the local linearizations are only approximations to the embedded curves, which in turn are imperfect representations of the high dimensional ground truth. This suggests that a multivariate characterization of the dynamics in terms of replicable patterns, i.e. clusters, of concurrent multivariate EVOdFNC RI would induce *meta*-EVOdFNCs (**Figure 9**) that capture more of the high-dimensional data variability than the RI of individual EVOdFNCs. In practice, for the data evaluated in this study, the resulting weight-vector centroids (**Figure 13**, center right) were each highly concentrated on one EVOdFNC, yielding meta-EVOdFNCs that strongly resemble the dominantly-weighted EVOdFNC in the corresponding weight-vector cluster centroid (**Figure 13**, **Figure 10**). However, although focused on one dominant EVOdFNC, the meta-dFNCs include content from all EVOdFNCs which allows for more flexible representation of the data when warranted. Moreover, occupancy rate forces a harder segmentation than representational importance, yielding slightly different results. Delusions, for example, show positive effects on the RI of EVOdFNC 3 and on the occupancy rate of meta-EVOdFNC 2 (**Figure 14**) which is heavily weighted toward EVOdFNC 3. Delusions however also exhibit negative effects on the occupancy rate of meta-EVOdFNC 4, which is heavily weighted on EVOdFNC 7 whose RI is not significantly affected by Delusions.

**Figure 13.**
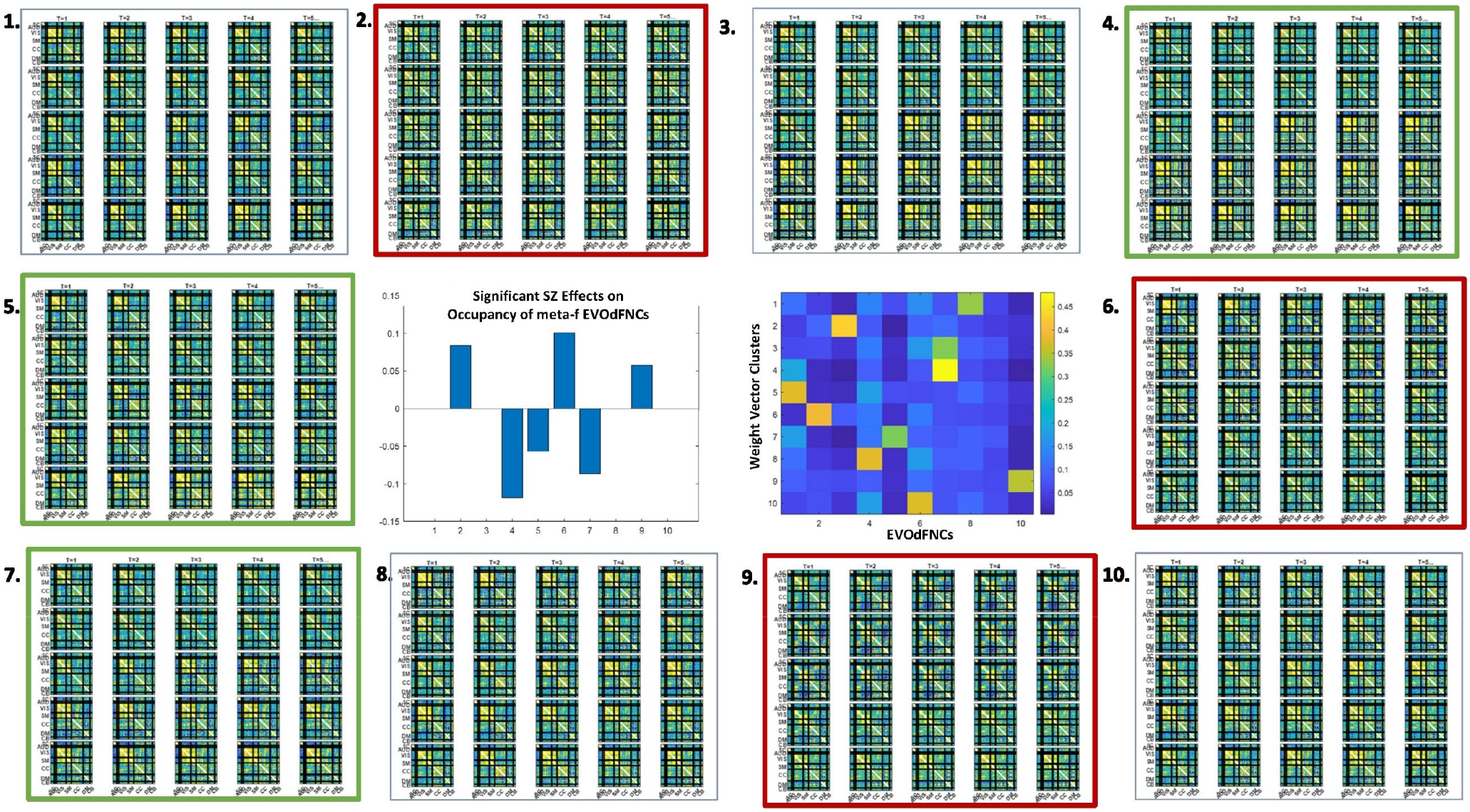
As with representational importance of EVOdFNCs, we see many significant group differences between schizophrenia patients and controls in the occupancy rates of meta-EVOdFNCs; thick red (resp. thick green) boxes designate significant positive (resp. negative) effect of SZ on meta-EVOdFNC occupancy rate; all displayed effects are significant at the *p* < 0.05 level after correction for multiple comparisons. The cluster centroids of the EVOdFNC weight vectors from which the meta-EVOdFNCs (middle row, column 3) are induced tend to focus the majority of weight on one EVOdFNC, so there is evident resemblance between meta-EVOdFNCs and the EVOdFNCs. Note that positive symptoms exhibit no significant (p<0.05, uncorrected) effects on the occupancy rates of the SNAPdFNCs.

**Figure 14.**
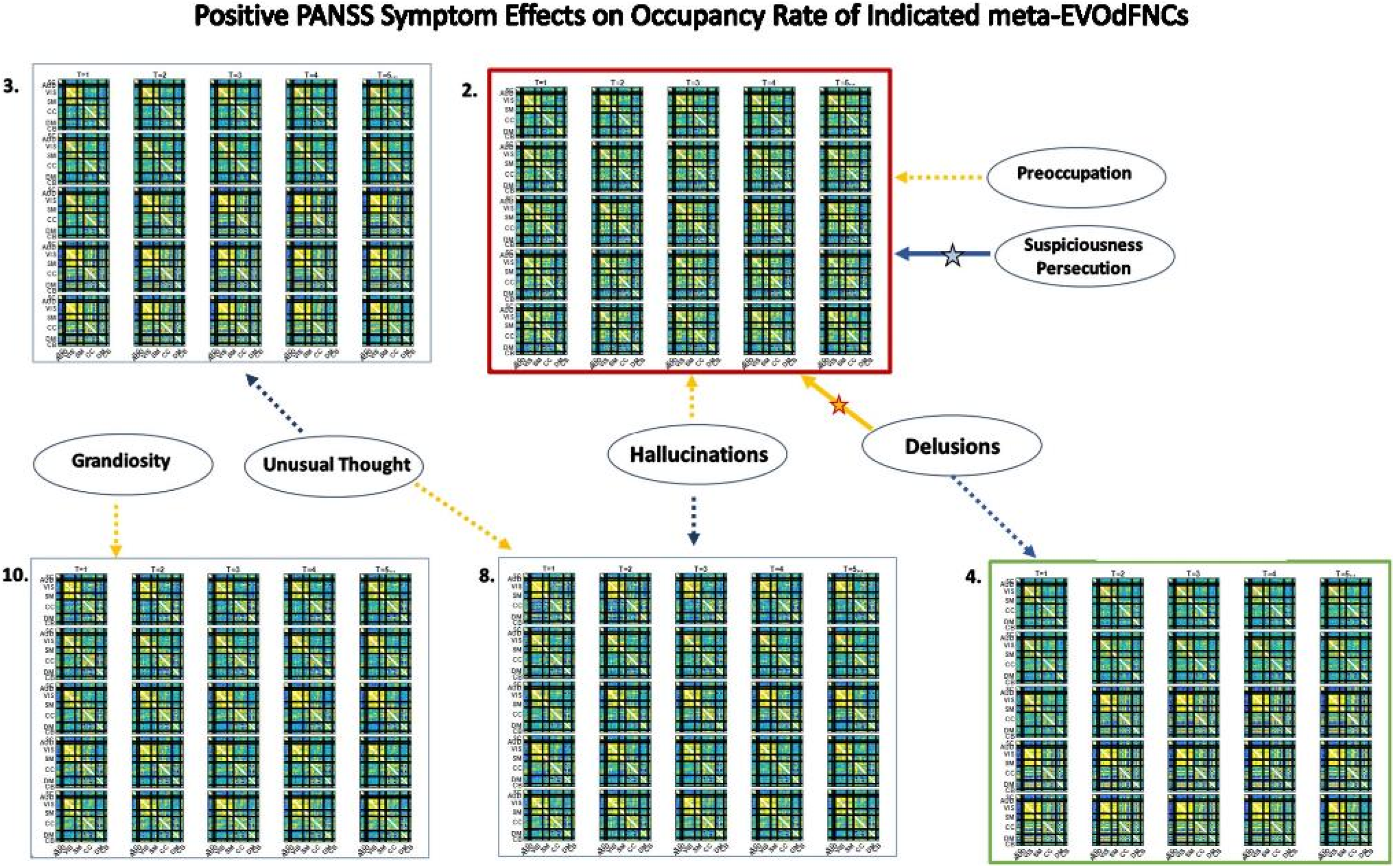
Significant positive symptom effects on the occupancy of meta-EVOdFNCs, each a weighted combination of EVOdFNCs. Thick arrows with stars indicate effects that are significant after correction for multiple comparisons; thin dashed arrows indicate effects that are significant with *p* < 0.05, but not after correction for multiple comparisons. Yellow lines denote positive effects. Blue lines denote negative effects. Consistent with results on representational importance of EVOdFNCs, meta-EVOdFNC 2 which is heavily weighted on EVOdFNC 3 is a locus of positive symptom effects,, including positive effects of Delusions and negative effects of Suspiciousness/Persecution that remain significant after correction for multiple comparisons. Note that positive symptoms exhibit no significant (p<0.05, uncorrected) effects on the occupancy rates of the SNAPdFNCs.

### 3. Occupancy Rates of Underlying 2D Linear Exemplar Clusters: Effects of Positive Symptoms and Relationship to EVOdFNCs

It is also possible to explore diagnosis and symptom effects directly in the 2D embedding, considering the linear exemplar cluster membership of each local linearization to an embedded subject trajectory (**Figure 15**). This approach “trusts” the embedding to distill some important relationships in the data while unavoidably distorting others. Surprisingly, there are no significant SZ effects on exemplar cluster membership of 2D local linearizations: nothing significant at the *p* < 0.05 level after correction for multiple comparisons, nor any raw uncorrected p-values less than 0.05. However, in the case of positive PANSS symptoms we find a number of significant relationships between positive PANSS scores and linear exemplar cluster membership (**Figure 16**), one of which, the negative effect of Hallucinations on the linear exemplar that lifts to EVOdFNC 8, is significant at the *p* < 0.05 level after correction for multiple comparisons.

**Figure 15.**
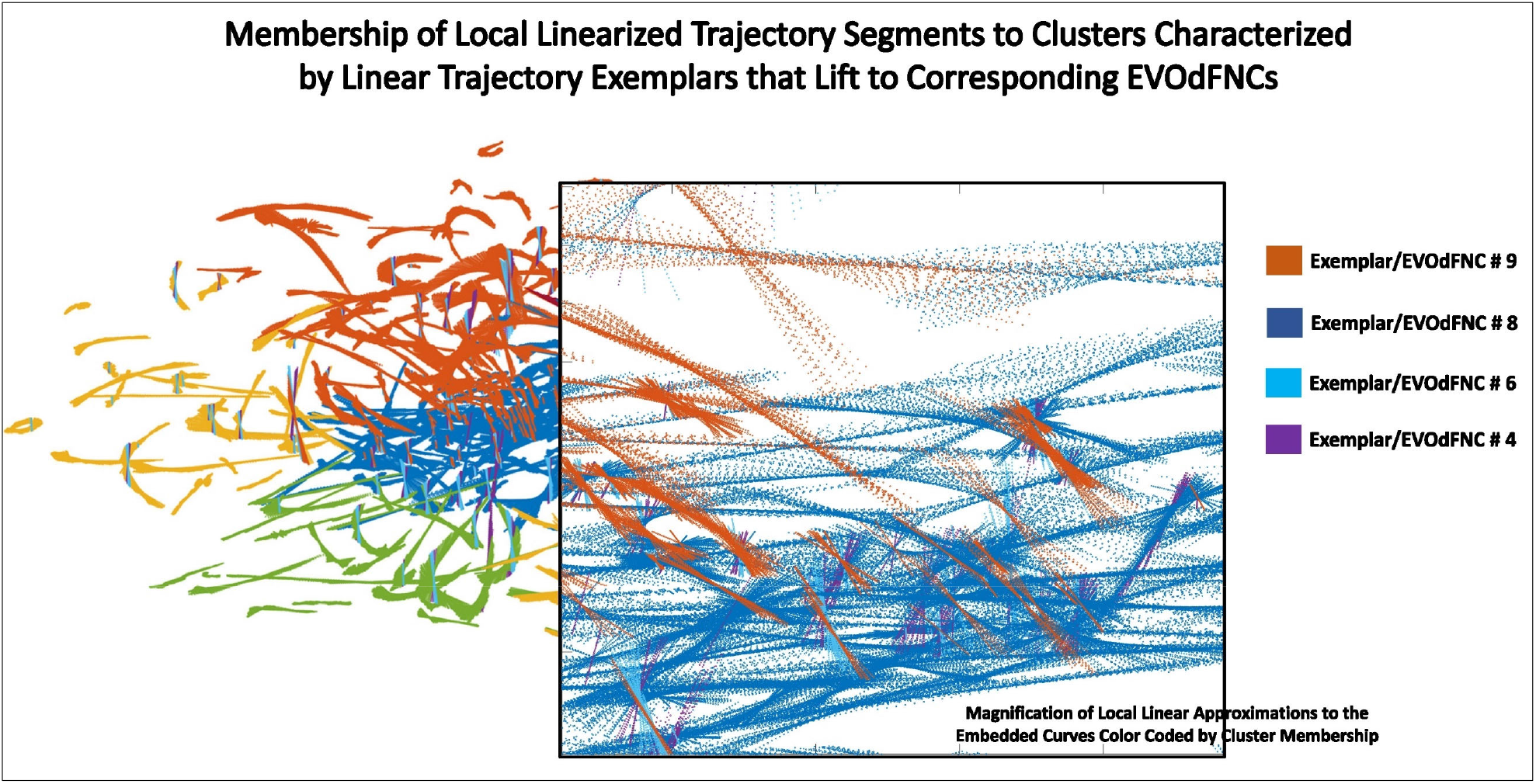
Zooming in on a box within the display (transparent background) of local linearizations of embedded curves (LTSs) colored according to the 2D linear trajectory exemplar cluster to which they belong. Within the same region the individual LTSs can belong to different exemplar clusters based on slope.

**Figure 16.**
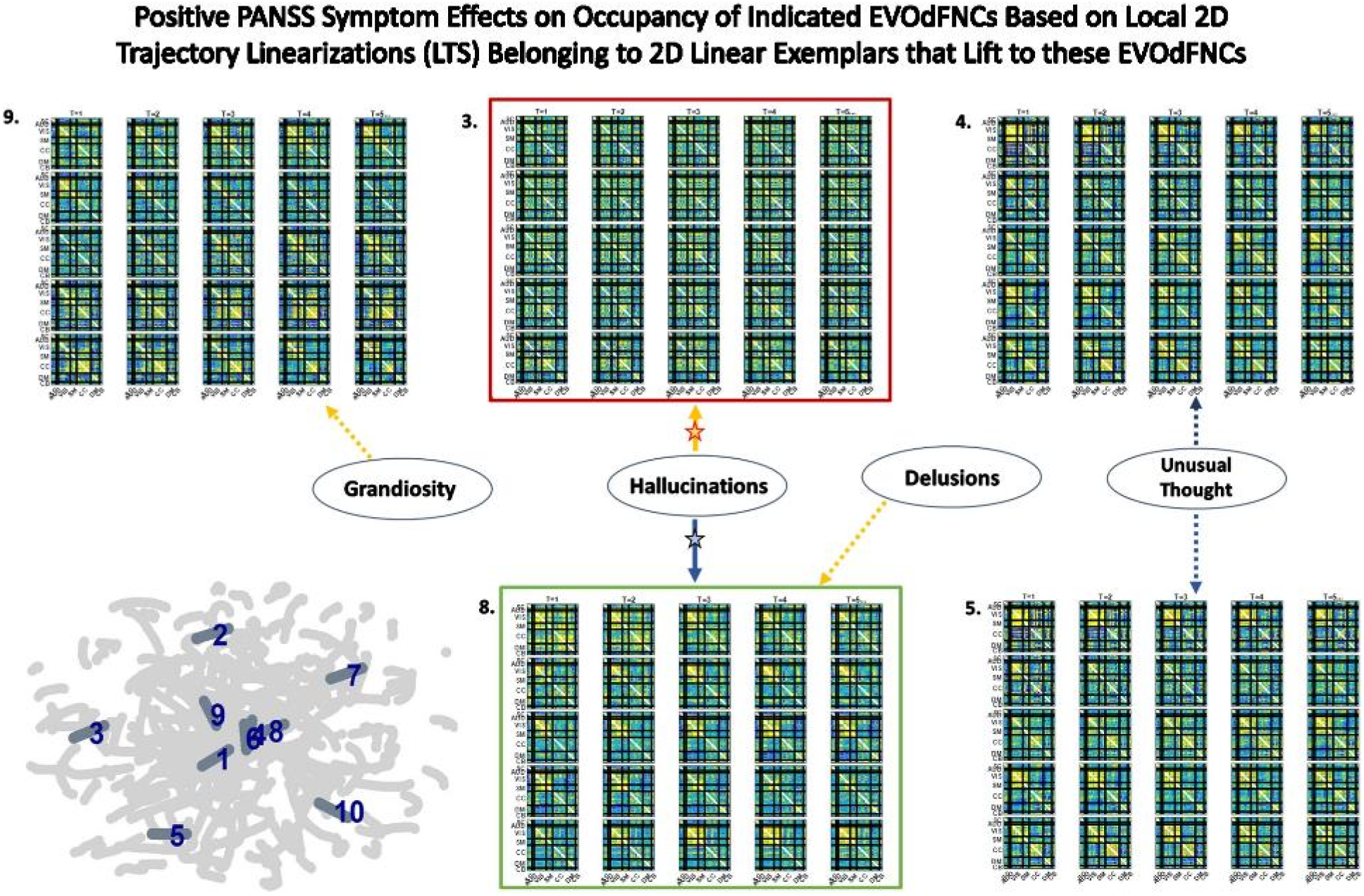
Five of the seven positive PANSS symptoms exhibited significant relationships with membership of local 2D trajectory linearizations to clusters defining the linear trajectory exemplars that lift to the displayed EVOdFNCs. Red (resp. green boxes around EVOdFNCs indicate positive (resp. negative) SZ effects on their representational importance in observed high-dimensional dFNC trajectories. Yellow (resp. blue) arrows point symptoms to EVOdFNCs with which they exhibit positive (resp. negative) membership effects. Symptom effects are significant at the *p* < 0.025 level. The negative effect of Hallucinations with 2D exemplar #8, i.e., with EVOdFNC #8, remains significant at the *p* < 0.05 level after correction for multiple comparisons. Note that positive symptoms exhibit no significant (p<0.05, uncorrected) effects on the occupancy rates of the SNAPdFNCs.

High-dimensional EVOdFNCs are inverses or lifts of 2D linear exemplars that summarize the planar embedding of group level high-dimensional dynamics. The exemplar membership of successive linear approximations to the embedded curves offers a “first pass” view of the alignment of subject connectivity dynamics with directional trends in the embedded data. As an embedding-intrinsic measure, exemplar cluster membership of 2D linear approximations are less concretely empirical than high-dimensional EVOdFNC representational importance. For positive SZ symptoms this embedding-intrinsic point of view strengthens the uncorrected significance of, e.g. the Hallucinations effects on EVOdFNCs 3 and 8 while identifying several new relationships between positive symptoms and dimensionally reduced dFNC dynamics (**Figure 16**).

## 4. DISCUSSION

Characterization and analysis of the time-varying resting state connectome continues to rely heavily on identifying a small number of fixed whole-brain connectivity patterns that manifest on timescales shorter than the full scan duration. The small set of fixed “states” is then then employed to model brain dynamics as a stationary Markov process, with the brain occupying and transitioning between this small set of fixed patterns. More sophisticated analyses of the complex dynamical processes that support human cognitive, emotional, executive and motor functions require frameworks for characterizing and leveraging the fluidly varying high dimensional dynamics presented by functional imaging modalities such as fMRI. Here we introduce an approach that works from a data-driven inversion of the summary gradients in a planar embedding of the high-dimensional dynamics (**Figure 17**) to capture group-level multiframe evolving “movie-style” representations of dynamic functional network connectivity (EVOdFNCs) in a large schizophrenia imaging study. We show that the method produces interpretable, naturalistic high-dimensional EVOdFNC states, whose contributions to HC and SZ dynamic connectivity differ significantly. The EVOdFNCs also expose distinct ways the two groups manifest and recede from certain characteristic organizational states of the connectome, e.g., the pattern in which DMN is anticorrelated with other networks and non-DMN networks are positively intercorrelated with each other (DMNneg).

**Figure 17.**
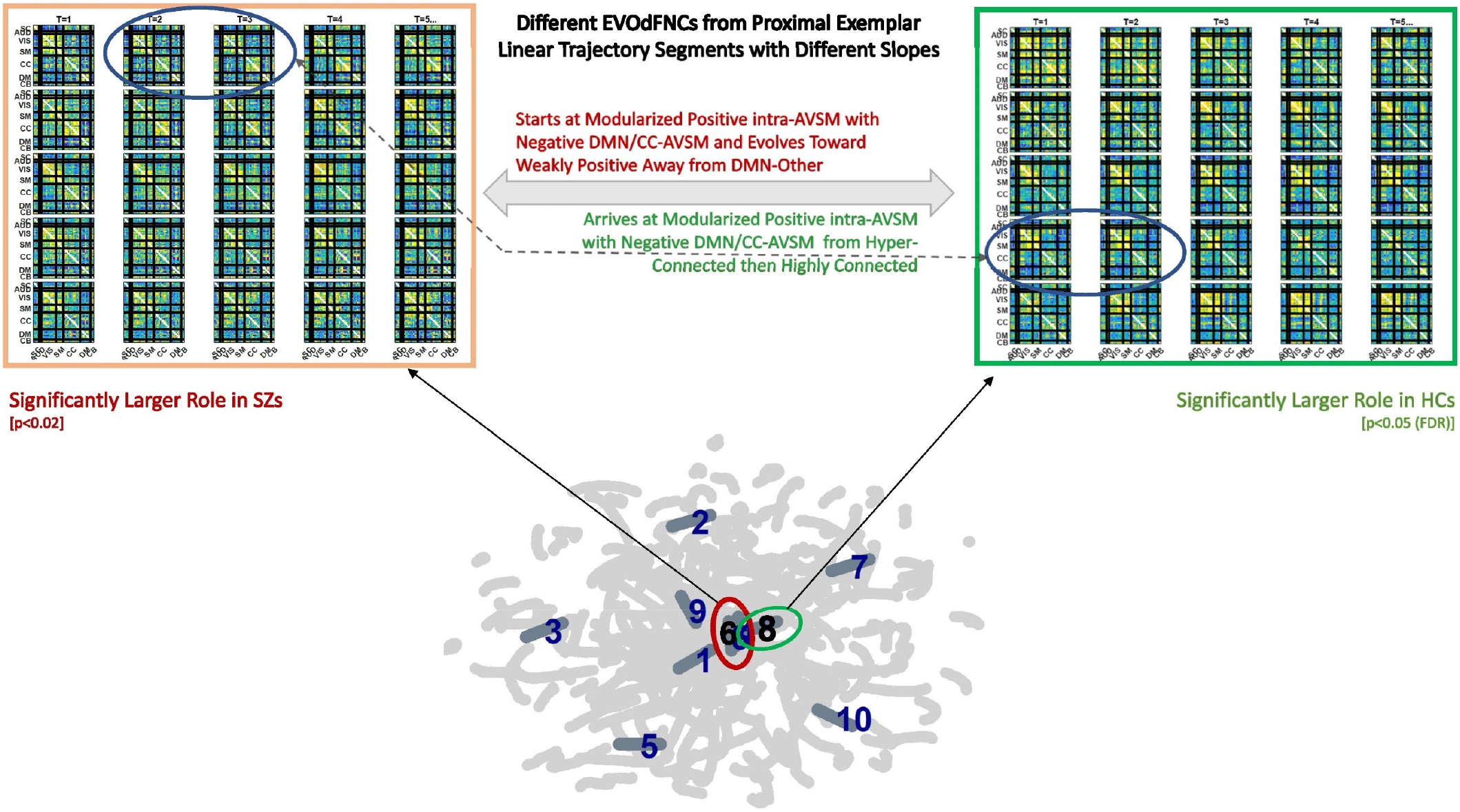
Top row shows two EVOdFNCs: leftmost in thick orange box is significantly (*p* < 0.025) more important in SZ before correction for multiple comparisons; the rightmost in the thick green box is significantly more important in HCs after correction for multiple comparisons. Note the similarity between the circled frames in both EVOdFNCs arising from the corresponding 2D linear trajectory exemplars that pass very near each other, but have different slopes; the EVOdFNC with higher representational importance in healthy controls corresponds to the more horizontal exemplar (#8 in the thick green circle) while the nearby exemplar (#6 in the thick red circle) lifts to EVOdFNC 6 which has significantly higher RI in SZs. Although the EVOdFNCs in the top row are built from 2D linear trajectory exemplars that are proximal, especially near their endpoints, their differing slopes (i.e., angles of approach) cause them to represent dynamic trajectories sufficiently different as to be to represent high dimensional dynamic contexts that are characteristic in one case of HCs and in the other of SZs.

Capturing characteristic longer sequences of evolving whole-brain connectivity exposes differences between patients and controls that reside in a complex joint feature space over temporal pacing, modular patterning and overall strength of functional integration. This richer viewpoint shows, e.g., that a DMNneg pattern with weak contrast and limited temporal variation (EVOdFNC 3) is highly implicated in both SZ and its positive symptoms, while high-contrast DMNneg dissolving into dysconnectivity (EVOdFNC 2) strongly differentiates SZ from HC but displays no sensitivity to positive symptoms. Finally, EVOdFNC 7 that, like EVOdFNC 2, starts with high-contrast DMNneg but instead of losing energy and modularity over time, gains connectivity strength while reconfiguring its modular patterning, is negatively correlated with SZ, while being both positively and negatively associated with positive SZ symptoms. Of the five EVOdFNCs that exhibit AVSM-dominant modularity, four (EVOdFNCs 1, 5, 7 and 8) are more representationally important in HCs and one (EVOdFNC 4) is highly important across the population (**Figure 19**) but not significantly different between SZs and HCs. At the timescale being examined, *τ* = 44 TRs (88 sec.), the majority of the EVOdFNCs captured pass through several discernibly different configurations. This is true for all of the EVOdFNCs with significantly greater representational importance in HCs (1, 5, 7 and 8) and for two of the EVOdFNCs with significantly greater RI in SZs (2 and 10). Of the more static EVOdFNCs (3 and 6) (see **Figure 18**), the one (EVOdFNC 3) that features consistent, low-contrast DMNneg plays a central role in positive symptom levels as well SZ generally; the other (EVOdFNC 6) is consistently diffusely disconnected and has not significant role either in distinguishing patients from controls, or in connection with positive symptoms. Intervals of diffuse dysconnectivity are associated with significant group differences and symptom effects when they appear in as subintervals in more EVOdFNCs (1, 2, 5, 10) that also have higher contrast intervals (see **Figure 18**). The role of disconnected periods in the evolving connectomes of HCs and SZ differ in timescale and their role as “connective tissue” between higher-magnitude patterns of connectivity, suggesting that clinically relevant aspects of resting-state connectivity dynamics are obscured by the field’s current overreliance on a simplifying Markov assumption. Fluidly evolving dynamic representations can also reveal important new connectivity patterns that are on the pathway from one familiar connectivity pattern from SNAPdFNC to another, e.g. the end of EVOdFNC 9 where intra-CC connectivity is stronger than AVSM connectivity and the negative connectivity between SM and part of the VIS with the rest of VIS in EVOdFNC 10.

**Figure 18.**
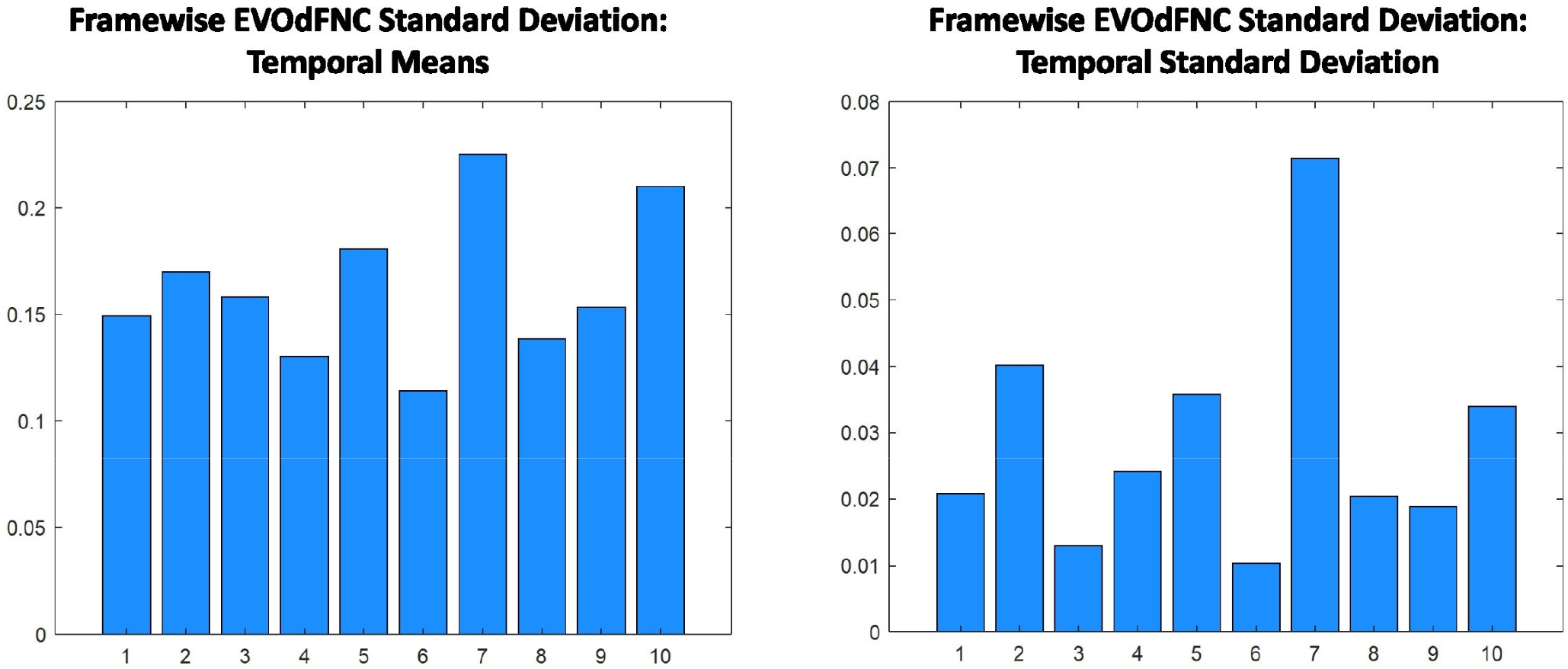
(Left) The average standard deviation per frame of each EVOdFNC (enumerated on the x-axis), a rough metric of how much functional contrast the EVOdFNC exhibits; (Right) The standard deviation through time of the amount of framewise functional contrast, i.e., the framewise standard deviation, in each EVOdFNC, roughly measuring the range of functional contrasts exhibited over the duration of the EVOdFNC.

It is also important to mention that, while promising, this approach has a number of limitations that will benefit from further development. The method has a large number of parameters, from those that govern the UMAP embedding to the temporal scale (duration of EVOdFNCs) to the number of clusters at both the 2D exemplar and meta-EVOdFNC stages. There are different ways of assessing representational importance, each of which has benefits and drawbacks. Finally, there are multiple levels of results, emphasizing either the embedding itself, each high-dimensional EVOdFNCs individually, or weighted combinations of the EVOdFNCs. In this study, each meta-EVOdFNCs was strongly focused on one specific EVOdFNC, so results from the individual and multivariate levels tracked each other. The relationship between exemplar cluster membership, defined within the planar embedding, and the high-dimensional representational importance of EVOdFNCs to observed sequences of dFNCs is more complicated (**Figure 19**), leading to some shared results at a statistical level (e.g., significant effects of Hallucinations and Delusions on exemplars/EVOdFNCs 3 and 8) with weaker relationships on a timepoint-by-timepoint basis (**Figure 17**, **Figure 20**, **Figure 21**). While acknowledging limitations, we believe this is an important first step toward more sophisticated analysis of high dimensional functional imaging data, allowing researchers to more finely resolve the relationship of longer dynamic *processes* to human health and performance.

**Figure 19.**
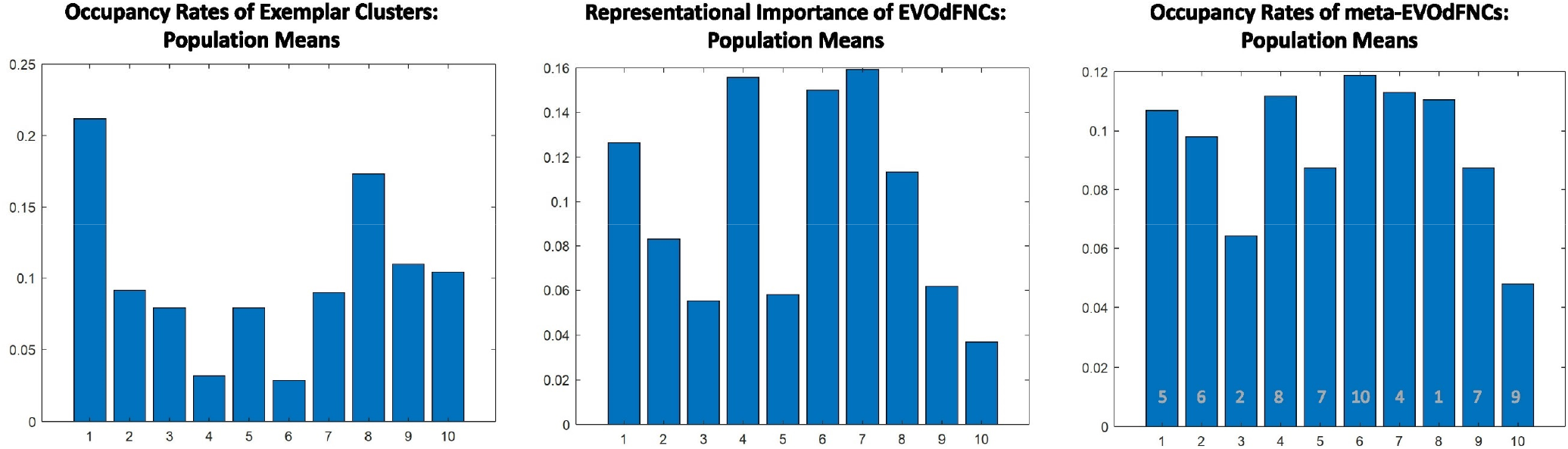
Population means of the occupancy rates of the linear exemplar clusters (Left), the representational importance of each EVOdFNC (Middle), and the occupancy rates of the meta-EVOdFNCs (Right); the x-axis in all panels tracks with the indices of linear exemplars, which map directly to the corresponding EVOdFNCs, but meta-EVOdFNCs are re-indexed to appear in the same order as the exemplar/EVOdFNC that the meta-EVOdFNC (with original index shown in gray) is most heavily weighted on (see **Figure 13**, middle row, 3^rd^ from right). The occupancy rate of 2D exemplar clusters is weakly negatively correlated (ρ = –0.115 (*p* = 0.7)) with the representational importance of the corresponding EVOdFNC. In particular, we see that 2D exemplars 4 and 6 are least occupied, but the EVOdFNCs they induce are empirically very important in representing high dimensional data. The occupancy rates of meta-EVOdFNCs is, however, highly correlated ((ρ = 0.89 (*p* < 0.0001)) with the representational importance of the EVOdFNCs they are most heavily weighted on.

**Figure 20.**
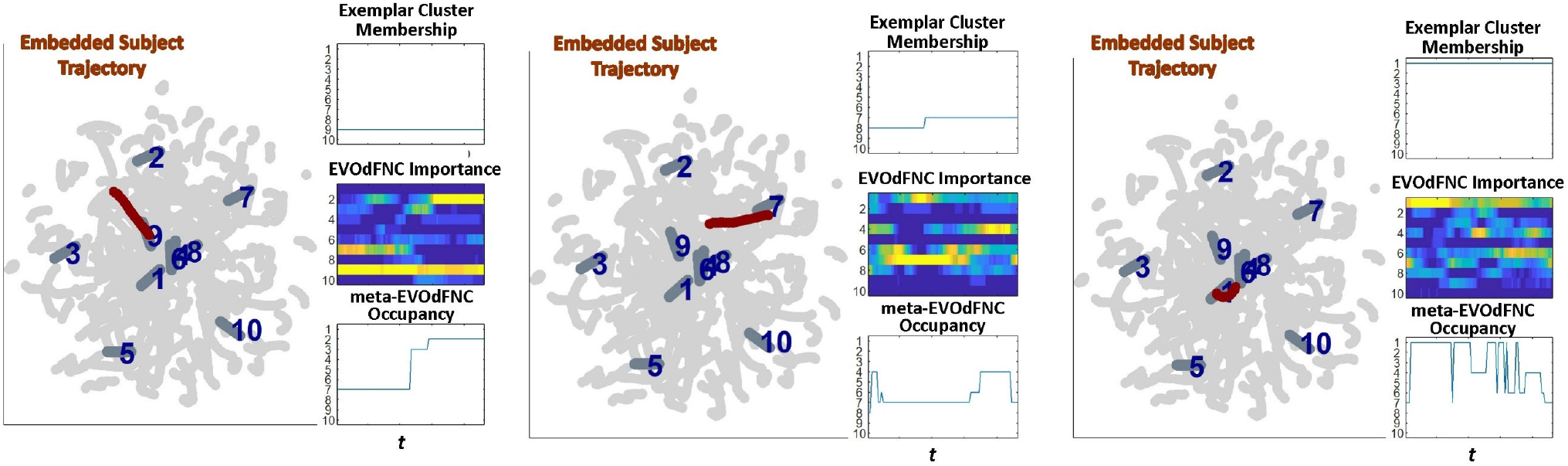
Three examples (leftmost panel in each subfigure) showing how proximity of subject embedded trajectories 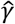 (maroon, superimposed with the ten linear exemplars (dark gray, labeled in dark blue) over the full embedding (light gray)) are represented in terms of (right panel in each subfigure, top row) membership of local linear approximations to 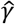 in clusters defining each of the ten linear exemplars; (right panel in each subfigure, middle row) the representational importance of the high-dimensional EVOdFNCs corresponding to each 2D linear exemplar in the subject’s high dimensional dFNC trajectory Γ, and (right panel in each subfigure, bottom row) the occupancy of meta-EVOdFNCs re-indexed to correspond with the EVOdFNC which is most strongly weighted in each meta-EVOdFNC

**Figure 21.**
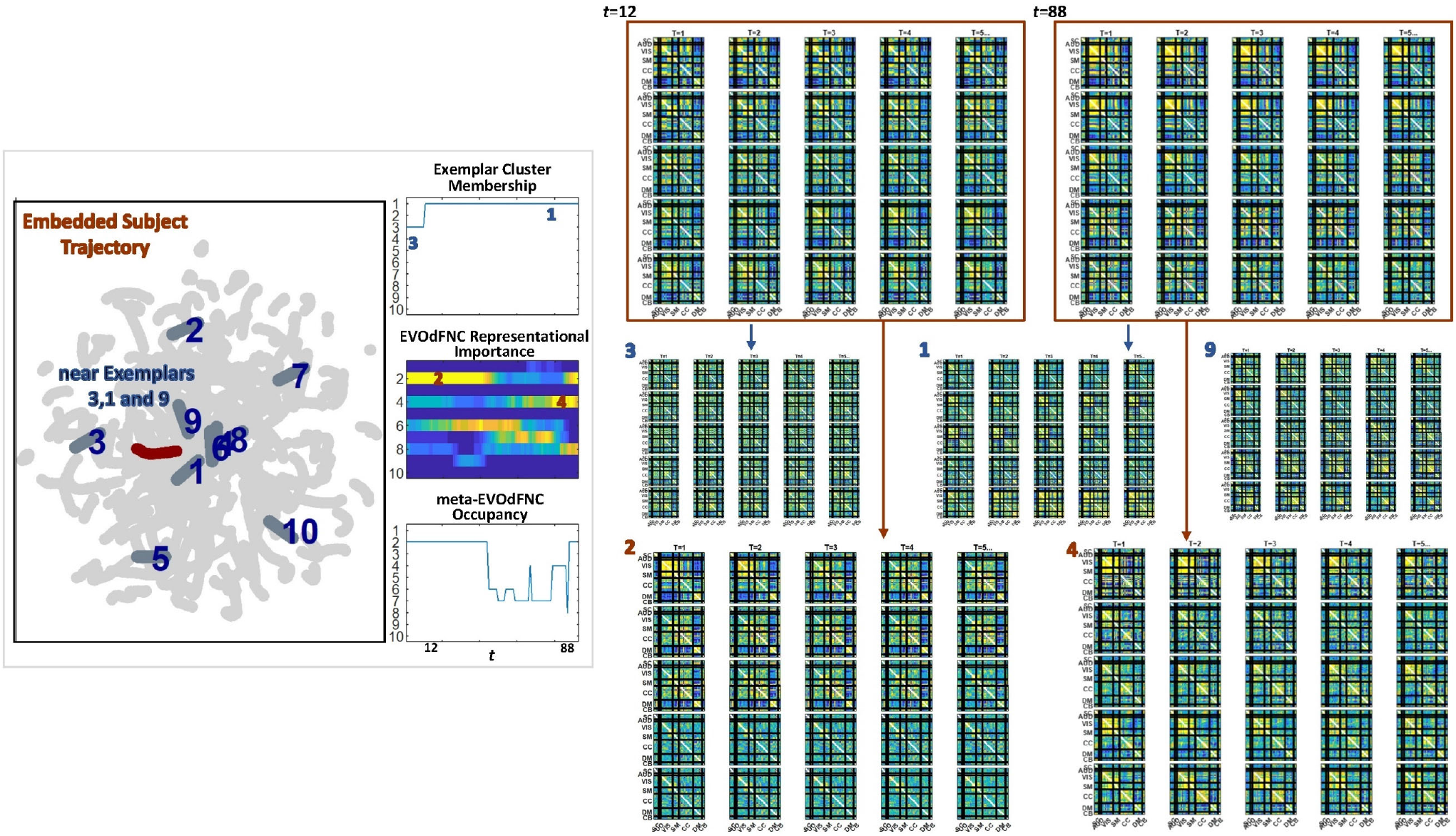
(Leftmost panel) One subject’s embedded trajectory 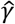 (maroon) and the ten linear examplars (dark gray, labeled in dark blue) shown superimposed over the full embedding (light gray). The embedded trajectory passes near exemplars 3, 1 and 9; (Left panel, column 2, top) As expected, the local linearizations approximating 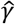 belong to clusters defining exemplar 3 and then exemplar 1, both of which 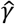 passes close to; (Left panel, column 2, middle); the time-varying representational importance of the high-dimensional EVOdFNCs corresponding to each linear exemplars in the subject’s high-dimensional dFNC trajectory Γ; the highest RI concentrates in EVOdFNCs 2 and 4 which are lifted from exemplars 2 and 4, neither of which is proximal to 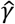; EVOdFNCs 3 and 1, corresponding to exemplars 3 and 1, have very low high-dimensional RI; (Right panel, top row) windows *w*_12_ and *w*_85_ starting at *t* = 12 and *t* = 85 which carry high RI from EVOdFNCs 2 and 4 respectively (Right panel, middle row) EVOdFNCs corresponding to exemplars 3,1 and 9 that 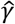 passes close to in the plane are not highly frame-wise correlated with Γ(*w*_12_) or Γ(*w*_85_) while (Right panel, bottom row) EVOdFNCs 2 and 4 which are most representationally important in Γ do exhibit evidently strong framewise correlations with Γ(*w*_12_) and Γ(*w*_85_).

## SUPPLEMENTARY MATERIAL

**Figure 22.**
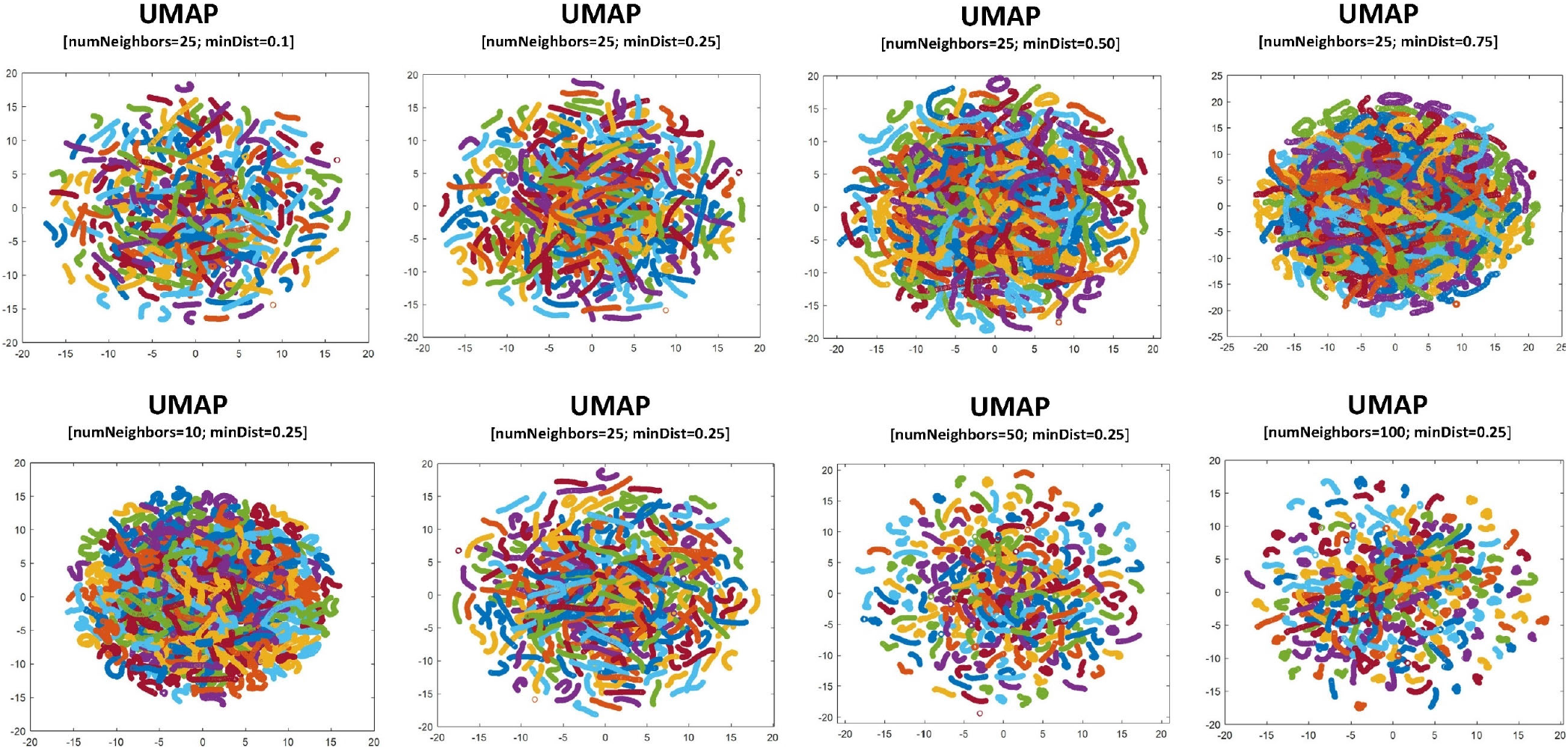
(Top Row) Holding the number of neighbors parameter constant, embedded trajectories are less tightly selfcontained as the minimum distance parameter grows. In a large dataset this has the effect of making the samples more densely intermingled; (Bottom Row) Holding the minimum distance parameter constant, the embedded trajectories shrink away from each other as number of neighbors parameter is increased.

**Figure 23.**
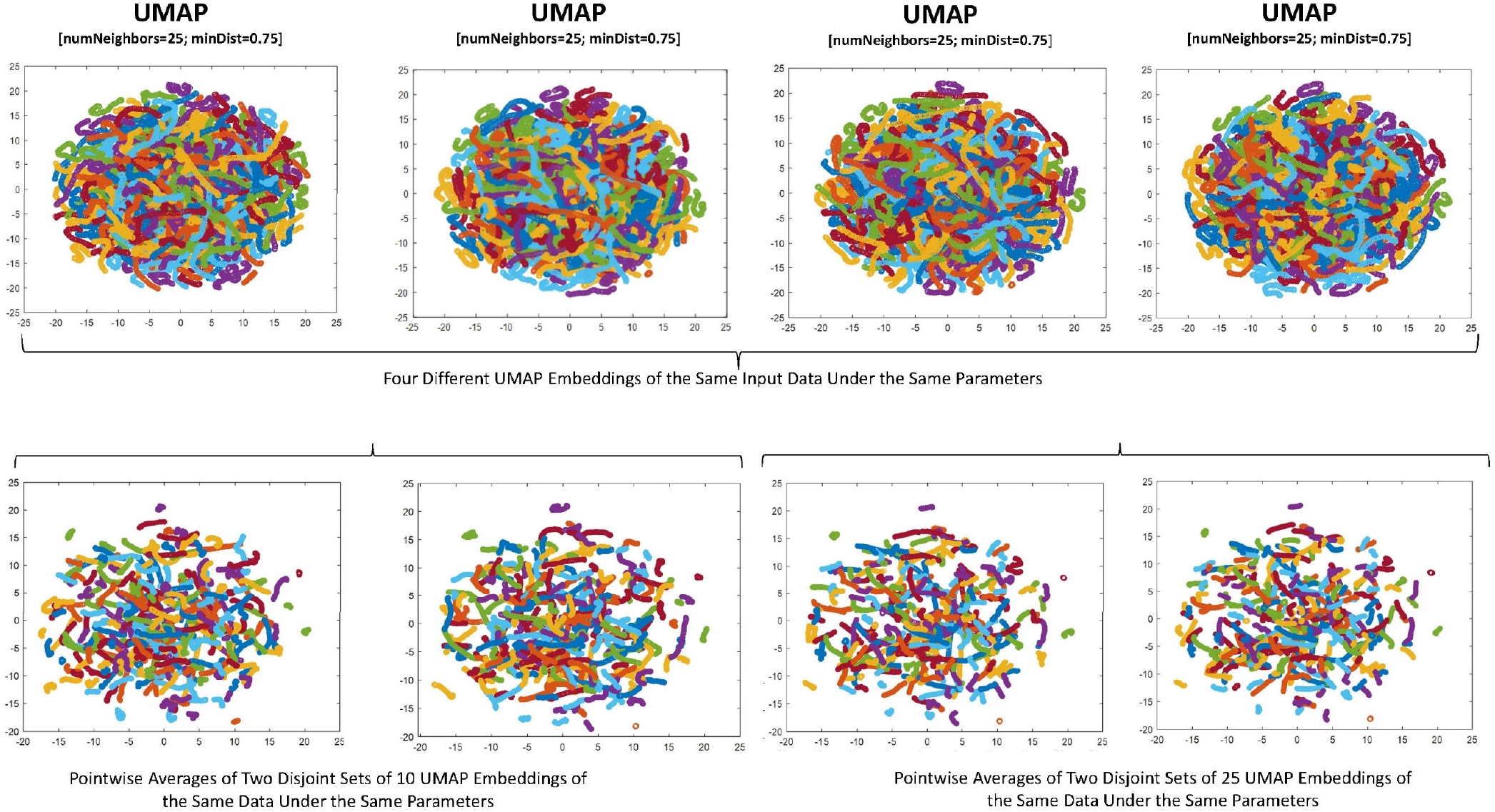
(Top Row) Four runs of UMAP on the data used in this study, at the parameter values used in this study. Each subject is represented in a different color. Many of the colors are visually indiscernible because the number of subjects is large. However, it is clear that subjects embed differently on different runs, though often occupying similar relative positions between runs; (Bottom Row, Columns 1 and 2) Averaging two disjoint sets of 10 different runs is slightly more stable than an arbitrary two individual runs; (Bottom Row, Columns 3 and 4) Averaging two disjoint sets of 25 different runs is considerably more stable than an arbitrary pair of individual runs. Note that the averaging distorts the qualitative properties of the embedding, creating more spatial separation between the averaged trajectory embeddings than is present in the individual runs. The high dimensional proximity of trajectories from different subjects is therefore less conserved in the averages. In spite of this distortion, we use an average of 25 runs for the work presented here due to the greater stability conferred by averaging.

**Figure 24.**
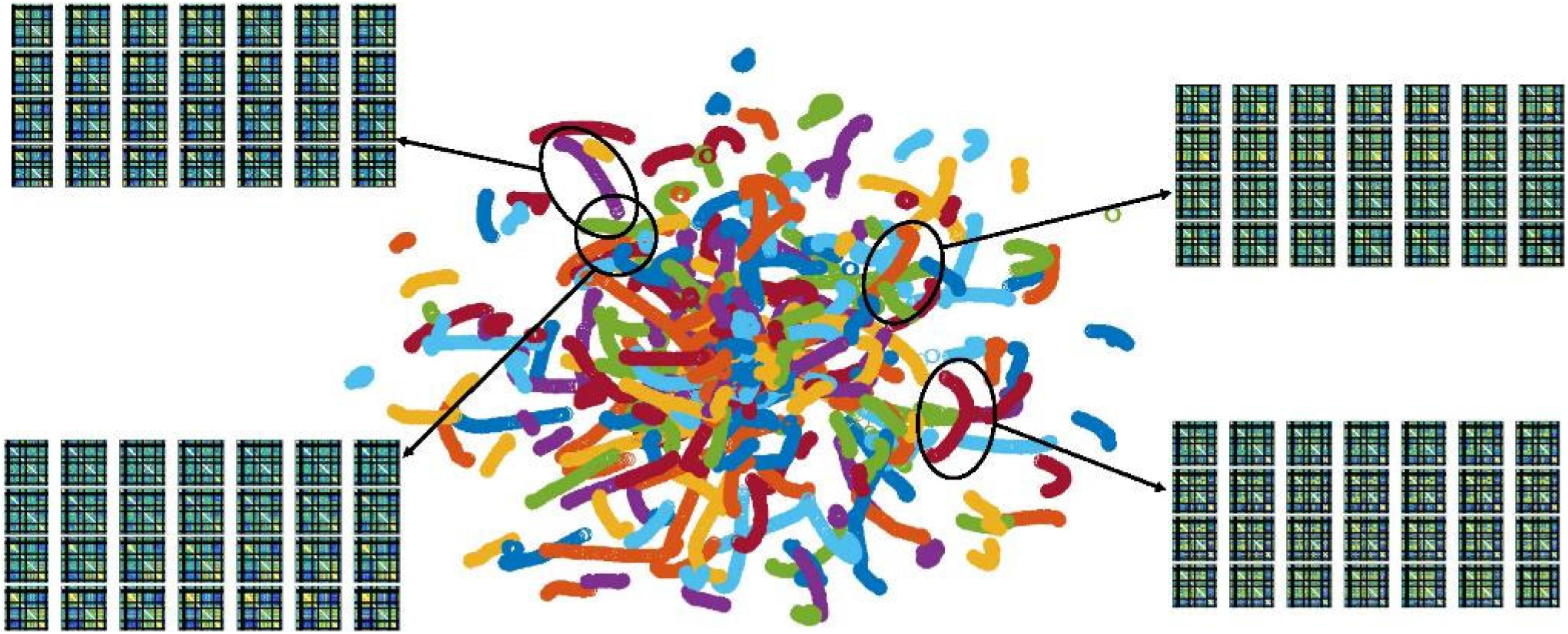
Although unsupervised, by preserving nearest-neighbor relations UMAP projects high dimensional continuous dFNC trajectories from each subject into contiguous temporally ordered curves in the plane (middle panel shows average over 25 UMAP runs; all observations (averaged) from a given subject share the same color coding; similar looking colors can be coding different subjects). Two pairs of proximal embedded trajectories are circled with arrows pointing to segments from the corresponding high-dimensional dFNC sequences illustrating the similarity of geometrically proximal embedded points and differences in high-dimensional sequencing for embedded trajectories in the same planar region.

